# Enhlink infers distal and context-specific enhancer-promoter linkages

**DOI:** 10.1101/2023.05.11.540453

**Authors:** Olivier B. Poirion, Wulin Zuo, Catrina Spruce, Sandra L. Daigle, Ashley Olson, Daniel A. Skelly, Elissa J. Chesler, Christopher L. Baker, Brian S. White

## Abstract

Enhancers play a crucial role in regulating gene expression and their functional status can be queried with cell type precision using using single-cell (sc)ATAC-seq. To facilitate analysis of such data, we developed Enhlink, a novel computational approach that leverages single -cell signals to infer linkages between regulatory DNA sequences, such as enhancers and promoters. Enhlink uses an ensemble strategy that integrates cell-level technical covariates to control for batch effects and biological covariates to infer robust condition-specific links and their associated *p*-values. It can integrate simultaneous gene expression and chromatin accessibility measurements of individual cells profiled by multi-omic experiments for increased specificity. We evaluated Enhlink using simulated and real scATAC-seq data, including those paired with physical enhancer-promoter links enumerated by promoter capture Hi-C and with multi-omic scATAC-/RNA-seq data we generated from the mouse striatum. These examples demonstrated that our method outperforms popular alternative strategies. In conjunction with eQTL analysis, Enhlink revealed a putative super-enhancer regulating key cell type-specific markers of striatal neurons. Taken together, our analyses demonstrate that Enhlink is accurate, powerful, and provides features that can lead to novel biological insights.

## Introduction

Gene transcription is regulated by non-coding DNA elements, called enhancers. Each consists of dense clusters of recognition motifs for sequence- and cell type-specific transcription factors (TFs), which bind and subsequently recruit coregulators, chromatin remodelers and modifiers, and RNA polymerase II (Panigrahi and O’Malley, 2021). A single gene can be regulated by multiple enhancers, with the cell type-specific activity of its enhancers conferring spatiotemporal control over it (Heinz *et al*., 2015; Robson, Ringel and Mundlos, 2019). Enhancer disruption and its concomitant modulation of target gene expression are increasingly recognized as disease-causing mechanisms (Corradin and Scacheri, 2014; Claringbould and Zaugg, 2021). In complex diseases, >90% of single nucleotide polymorphisms (SNPs) identified by genome-wide association studies (GWAS) are in non-coding regions of the genome far from promoters and potentially within enhancers (Claringbould and Zaugg, 2021). The link between an enhancer and its target gene (or, equivalently, promoter) needs to be established, and is an ongoing challenge in the field.

Enhancer-promoter links can be directly detected with experimental techniques including Hi-C, (Lieberman-Aiden *et al*., 2009) (Schoenfelder *et al*., 2018). However, complex protocols, high cost, low resolution (Galitsyna and Gelfand, 2021) and their inability to detect interchromosomal interactions (Panigrahi and O’Malley, 2021) currently limit their applications.

As an alternative, Pliner and colleagues demonstrated how links can be inferred *computationally* by exploiting measurements of chromatin accessibility from single-cell ATAC-seq (scATAC-seq) data at a gene’s promoter and its active enhancers during transcription. The authors first inferred open chromatin regions (OCRs) from “peaks” of reads in scATAC-seq data, and applied a computational method, Cicero, to identify enhancer-promoter pairs with correlated peaks of chromatin accessibility (Pliner *et al*., 2018). Cicero handles the sparsity of scATAC-seq data by aggregating binary accessibility data from similar cells into counts, with related cells determined through similarities in a low-dimensional embedding. It reduces batch effects, principally arising from library size, by adjusting aggregated counts. Finally, it addresses the high dimension inherent in genome-wide discovery by inferring regularized covariance matrices describing accessibility peaks.

Following Cicero’s pioneering approach, linkage inference from scATAC-seq has become a popular strategy in various exploratory (Wang *et al*., 2020; ‘A multimodal cell census and atlas of the mammalian primary motor cortex’, 2021)(Li *et al*., 2021) and methodological studies (Kamimoto, Hoffmann and Morris, 2020). Several other recent methods can also be used to infer enhancer-gene links from scATAC-seq data. Signac (Stuart *et al*., 2021), ArchR (Granja *et al*., 2021), and SnapATAC (Fang *et al*., 2021) are comprehensive toolkits for scATAC-seq analysis that include linkage inference methods. ArchR computes the Pearson correlation between accessible regions represented in a low-dimensional embedding of aggregated cells and derives a *p*-value from it. Signac also computes Pearson correlation *p*-values, but instead does so using a random background of enhancers. Finally, SnapATAC fits univariate logistic regression models to chromatin accessibility using gene expression as features. SnapATAC and Signac are designed to associate enhancer accessibilities with gene expression from either a multi-omic dataset or a matching scRNA-seq dataset which is combined with the scATAC-seq using a label transfer procedure (Fang *et al*., 2021). In contrast, ArchR has the ability to process scATAC-seq alone or in combination with scRNA-seq. To summarize, these approaches leverage correlations between enhancers and promoter accessibility or gene expression at the single-cell level to deduce enhancer-gene links.

However, single-cell experiments have continued to grow in size and complexity since these methods were developed, leading to exquisite contextual specificity for inference on enhancer-promoter interactions, but also producing additional difficulties that are not adequately addressed by existing computational methods. For example, our recent murine study made use of a factorial and hierarchical experimental design to characterize the contribution of genetics, sex, and diet to cellular heterogeneity in two metabolism-related tissues (Poirion *et al.,* 2023), by performing large-scale, single-cell sequencing across technical batches. Existing methods can not directly model the impact of biological covariates, nor can cell-binning approaches control for technical covariates that differ across cells *within* a bin.

To address the challenges of studies with complex experimental designs, we developed Enhlink, a novel approach for inferring enhancer-promoter co-accessibility. It detects biological effects and controls technical effects by incorporating appropriate covariates into a nonlinear modeling framework involving single cells, rather than aggregates. It selects a parsimonious set of enhancers associated with a promoter to smooth the sparse representation of any individual enhancer while prioritizing those with the largest effect. To do so, Enhlink uses a random forest-like approach, where cell-level (binary) accessibilities of enhancers and biological and technical factors are features and the cell-level accessibility of a promoter is the response variable. If multi-omic chromatin accessibility and gene expression measurements are simultaneously available for each cell, Enhlink can further prioritize enhancers by associating them with the expression of the promoter’s target gene. Unlike existing methods, Enhlink has the ability to predict both proximal and distal enhancer-gene linkages and identify linkage specific to biological covariates, while also integrating a simulation workflow that utilizes experimentally validated enhancer-promoter signals to optionally estimate prediction accuracy.

Using simulation parameterized by experimentally-validated enhancer-promoter pairs, we show that Enhlink minimizes false positives and negatives relative to other approaches. We further demonstrate that Enhlink results are resilient to technical batches in our T2D study and that it has superior precision evaluated using enhancer-promoter interactions detected from paired promoter capture (PC)Hi-C data. Finally, we generated a multi-ome single-nuclei (sn)ATAC- and RNA-seq dataset from a study of sex and strain differences in the mouse striatum, a brain region involved in motivated learning and associated with acute and chronic effects of drug addiction (Lobo and Nestler, 2011; Yager *et al*., 2015). After identifying the two main neuron populations defined by their expression of dopamine receptor 1 (*Drd1*) or 2 (*Drd2*), we inferred neuron-specific enhancer-promoter links using both promoter accessibility and gene expression. We identified strong putative *cis* and *trans* regulatory regions amongst the two classes of neurons that we also intersected with a set of genetic variants. Notably, we identified several enhancers 500kb downstream of the *Drd1* promoter that were directly correlated with the regulation of multiple distal genes involved in the Drd1/Drd2 genetic program and may act as a super-enhancer. Enhlink should similarly enable discovery of enhancer-promoter co-accessibility in other complex scATAC-seq and multi-omic snATAC-/snRNA-seq datasets.

### Inferring biologically meaningful co-accessibilities from sc/snATAC-seq data

To assess enhancer-promoter co-accessibility inference from snATAC-seq data, we used the results of a previously published human heart study that generated snATAC-seq data and experimentally validated enhancer-promoter pairs (Hocker *et al*., 2021). We focused on the *KCNH2* promoter, for which the study identified an enhancer with a nearby risk variant (rs7789146) (**Figure 1A**) linked to atrial fibrillation. That study validated the role of this variant on enhancer function via CRISPR-Cas9 genome editing of a human pluripotent stem cell-derived cardiomyocyte (CM) cell line (Hocker *et al*., 2021). Here, we determined whether the accessibilities of the rs7789146 enhancer and *KCNH2* promoter correlated across cells, by representing each as a Boolean vector whose entries reflect whether at least one snATAC-seq read was present within the respective region and cell. We then computed the accuracy, recall, and f1-score between the promoter and the enhancer vectors, separately for atrial (aCM) and ventricular (vCM) cardiomyocytes, and compared them to null score distributions (**Figures 1B and S1**; see Methods). We expanded this analysis to the promoter of *MYL2* and three of its putative enhancers, also highlighted by the original study (Hocker *et al*., 2021) (**Figure S1**). In all cases, the f1-score and recall values were significantly higher than random (*p* < 0.05; **Figures 1B and S1**). These results further justify computational methods that infer enhancer-promoter links from their co-accessibility in snATAC-seq data, while the enhancers associated with the *KCNH2* and *MYL2* promoters provide a means of evaluating such methods.

**Figure 1.**
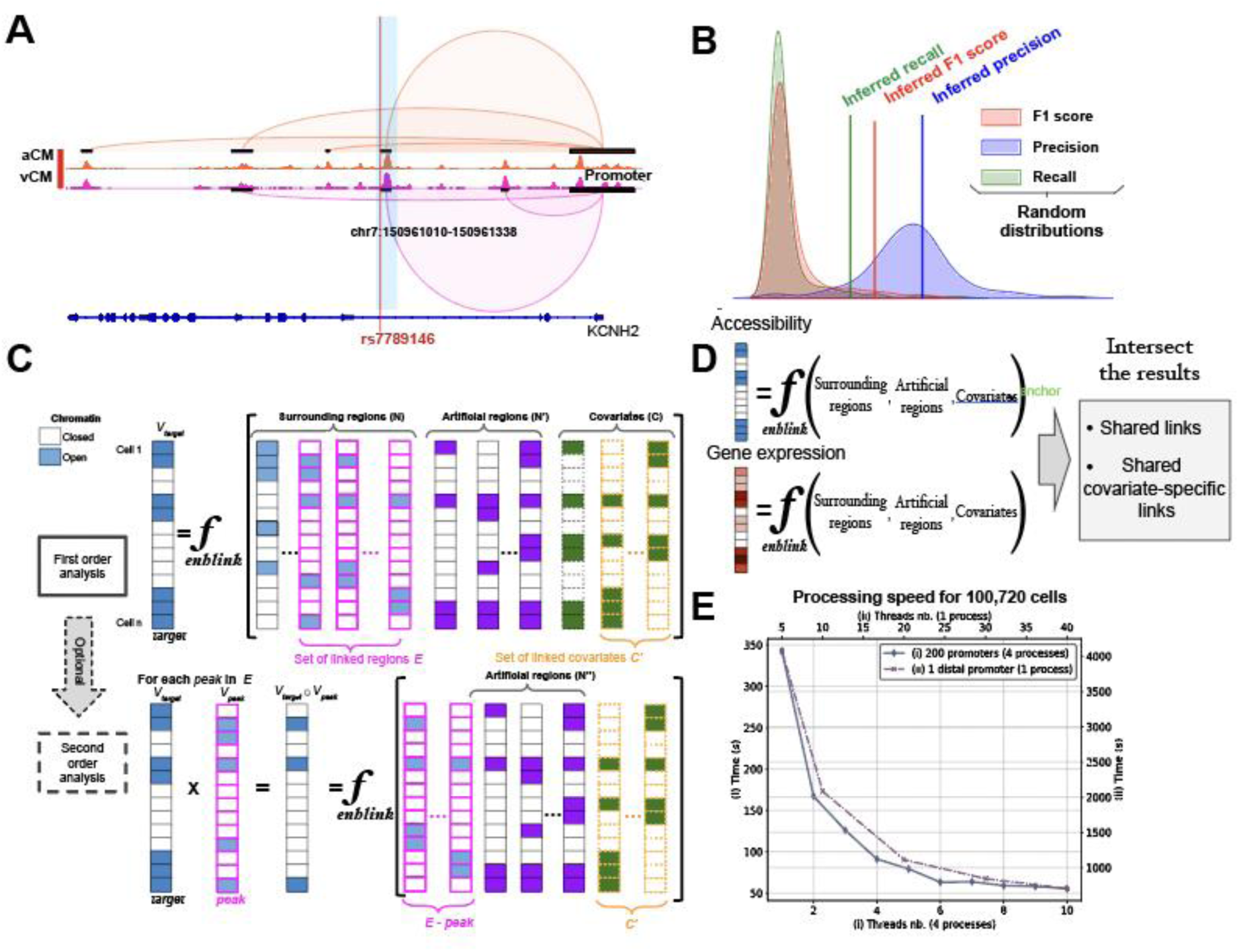
Enhlink infers linkage by modeling covariates, clusters and the surrounding enhancers. **A** Chromatin accessibility tracks with enhancer-promoter co-accessibility links inferred with Enhlink from human atrial (aCM) and ventricular (vCM) cardiomyocytes. The enhancer highlighted in blue was previously experimentally validated. **B** Accuracy scores computed from validated vCM enhancer/promoter pair for the promoter of *KCNH2* using scATAC-seq data and compared to their random score distribution. **C** Enhlink models a target region as a function of its surrounding genomic regions (i.e., enhancers) and biological and technical covariates. Artificial regions are added to reach a sufficient number of variables for computing feature scores and *p*-values. Enhlink can optionally perform a second-order analysis to identify covariates associated with links. **D** Enhlink can leverage multi-omics datasets by modelling a target region by either its accessibility or its expression and by intersecting the two resulting sets to identify links shared across both modalities. **E** Processing time for detecting associations (scenario I) for 200 promoters and their cis (+/- 250kb) OCR features from the islet dataset using four processes and (scenario II) between one promoter and 260,344 *cis* and *trans* OCR features using one process. Processing time (left axis for I and right for II) as a function of number of threads per process (bottom axis for I and top for II).

### Multi-omic inference of condition-specific enhancer-promoter links with Enhlink

Enhlink is a new approach that has been designed as an efficient computational framework for inferring co-accessibilities between OCRs, such as enhancers and promoters, from snATAC-seq data that are robust to technical batch effects. Enhlink parsimoniously identifies enhancer-promoter links across genome-wide candidates and can further prioritize associations that are supported by paired (multi-omic) expression data, where available (**Figures 1C and S2**). Enlink identifies OCRs and biological factors that “explain” a promoter, independent of technical factors (see Methods). It does so by using single-cell representations of features – promoters, enhancers, and biological and technical factors – where each is a binary vector with an element corresponding to the accessibility (or factor label) of each cell. The candidate set of enhancers may be limited to a genomic range surrounding the promoter (+/- 250kb, by default), to approximate the promoter’s topologically associating domain (TAD) – i.e., a three-dimensional subregion of the genome that sequesters self-interacting regions (McArthur and Capra, 2021), or may include all peaks genome-wide to model distal (Panigrahi and O’Malley, 2021) and indirect links. Biological and technical factors such as batch, lineage, and genotype, and categorical factors can be represented in full generality through “one-hot encoding.” Enhlink uses a binary decision tree to iteratively select features that maximize a modified information gain (see Methods). It computes an ensemble of such trees, bootstrapping the cells and selecting a random subset of features in each tree, in a manner similar to random forests. Bootstrapping accounts for heterogeneity across datasets and enables calculation of *p*-values for each enhancer or biological factor. The depth of each tree is controlled by an intuitive hyperparameter, which effectively sets the expected number of enhancers per promoter (four, by default). This depth and random feature subsetting prioritize a reduced set of enhancers (or biological factors) having the strongest, independent association with the promoter.

**Figure 2.**
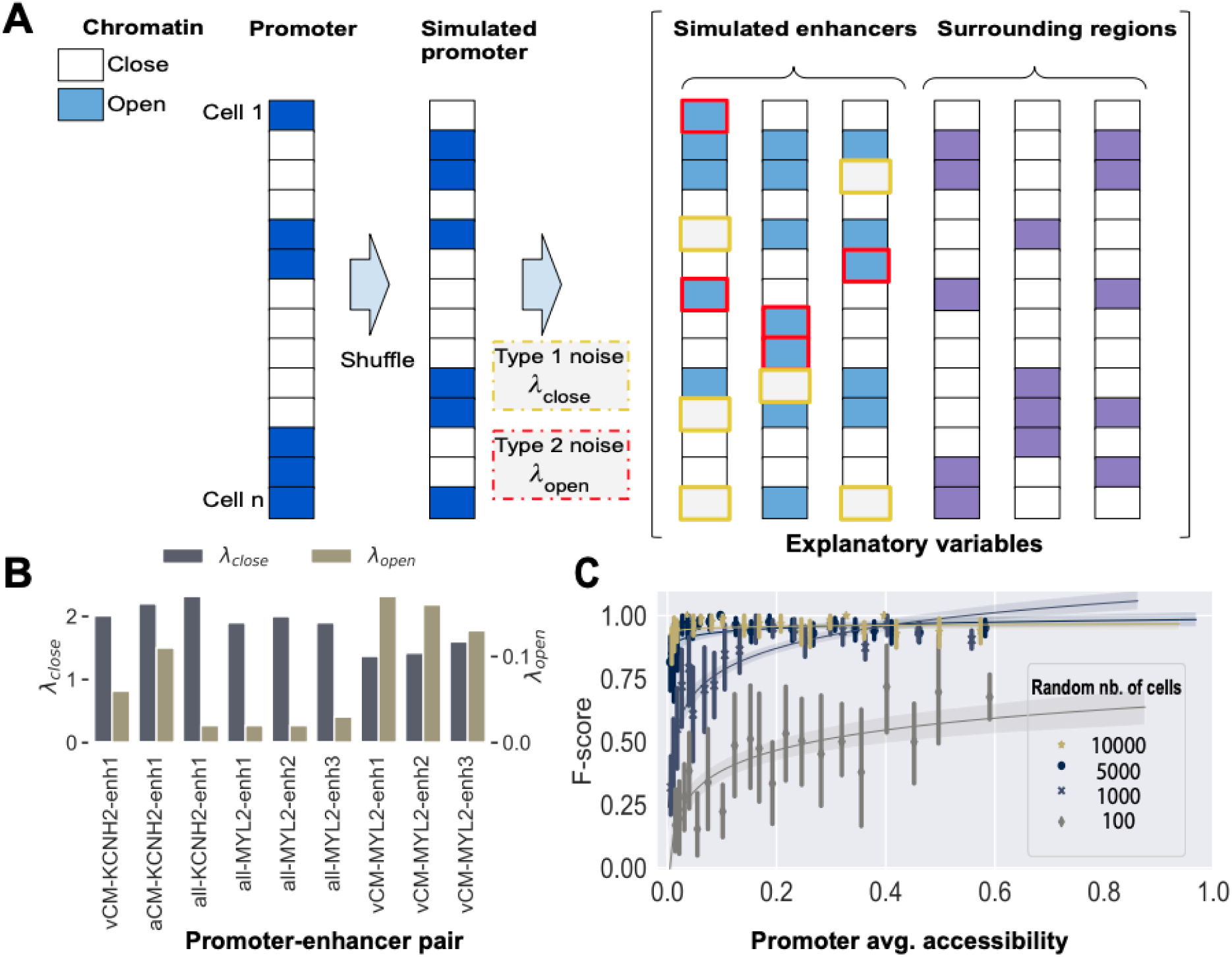
Empirically-parameterized simulation demonstrates Enhlink’s high accuracy. **A** Workflow to simulate promoter-enhancer associations parameterized by experimental data. The accessibilities of a promoter and its associated enhancers across cells are simulated from a single promoter-enhancer pair having a validated association. The simulated promoter accessibilities are derived by randomly shuffling the binary, scATAC-seq-derived accessibilities of the validated promoter across cells. Each simulated enhancer accessibility for a given cell is generated from the simulated promoter accessibility for that cell via a process that probabilistically flips the cell’s chromatin state: from closed to open (parameterized by λ_open_) or from open to closed (λ_close_). λ_open_ and λ_close_ are determined from the validated promoter-enhancer pair. The simulated enhancers are then integrated with the surrounding regions used as background. **B** λ_open_ and λ_close_ distribution parameters inferred from chromatin accessibility of enhancer-promoter pairs previously validated in human scATAC-seq cardiomyocyte cells (Hocker et. al 2021). Pairs involve the promoter *KCNH2* or *MYL2* as determined in all cells or in the subset of aCM or vCM cells. **C** f1-score (y axis) of simulated promoter-enhancer pairs as a function of average promoter accessibility and number of cells. Error bars summarize 20 simulated promoters. Each simulated promoter has between two and seven associated simulated enhancers.

Enhlink can further prioritize these snATAC-seq-derived enhancers by integrating mRNA measurements simultaneously assayed along with chromatin accessibility on each cell (Vandereyken *et al*., 2023). When such data are available, Enhlink identifies enhancers using both the promoter accessibility profiles and their associated gene expressions. By retaining enhancers that are concordant in both modalities, Enhlink enhances the likelihood of association between the identified enhancers and their target genes (**Figure 1D**). Enhlink then refines the snATAC-seq-derived set of promoter-associated enhancers by intersecting them with those derived from snRNA-seq.

Finally, Enhlink can identify enhancers active in a context-specific manner, e.g., those associated with a promoter in a specific biological condition or those cooperating with another promoter-linked enhancer. This is done via a second-order analysis in which the intersection (i.e., product of binary vectors) of a promoter and a biological factor (in the first case) or of a promoter and enhancer (in the second) are substituted for the accessibility profile of a promoter in the above framework (**Figure 1C**).

### Enhlink implementation and speed

Enhlink achieves its computational efficiency through its implementation in Go (https://go.dev/), a programming language optimized for CPU and memory usage (Dymora and Paszkiewicz, 2020). It computes each decision tree within a distinct thread, allowing computational speed to scale with the number of threads used (**Figure 1E**). Additionally, it can distribute the computation of a set of promoters or a grid of hyperparameters over multiple “processes”, improving the computational time while preserving the amount of memory needed on High-Performance Computing (HPC) clusters. Enhlink processed 200 promoters over 100,720 cells in 55 seconds using ten threads in each of four processes on a cluster of 52 CPUs and in 167 seconds with two threads in each of four processes (**Figure 1E**). Analysis of a single promoter using all 295,089 genome-wide peaks took approximately 700 seconds **(Figure 1E).** These processing times could have been further reduced with little impact on performance by randomly downsampling cells or peaks (see results section below). Memory usage only increased marginally with the number of threads and processes. Overall, Enhlink’s execution time for each promoter is linearly proportional to the number of trees in the ensemble, the number of enhancer peaks considered as features, and the number of cells, while it is exponentially dependent on the depth of each tree, and inversely proportional to the total number of threads used (across all processes). Enhlink takes as input sparse matrices in an MTX format compatible with Cell Ranger and easily generated from Python R workflows. Enhlink open-source code is freely available and accompanied by in-depth tutorials (https://gitlab.com/Grouumf/enhlinktools).

### Estimating accuracy and power analysis from simulated data

Inspired by the signal detected from snATAC-seq data for observed enhancer-promoter interactions (**Figure 1B**), we designed a strategy to simulate enhancer-promoter co-accessibilities with characteristics similar to those previously validated or well characterized. Briefly, we simulated a promoter accessibility vector by randomly shuffling one of the *MYL2* or *KCNH2* promoters observed within vCMs or aCMs and described above. We then simulated enhancer vectors by introducing random noise into the simulated promoter vector (**Figure 2A**). To model noise in the simulated enhancers, we defined the probability of a cell having a read at the promoter and not at the enhancer (λ_close_) or, conversely, at the enhancer and not at the promoter (λ_open_; see Methods). We estimated λ_close_ and λ_open_ from the observed, cell type-specific *MYL2* and *KCNH2* enhancers and found similar values for λ_close_ (1.9 +/- 0.2) and λ_open_ (0.08 +/- 0.3) (**Figure 2B**). We extended this analysis using all the cells and found slightly higher λ_close_ (2.0 +/- 0.2) and lower λ_open_ (0.02 +/-0.05) values. We then estimated Enhlink precision, recall, and f1-score using the simulated enhancer-promoter associations as ground truth. Simulation across three datasets, for a large number of promoters, and with multiple λ_close_ and λ_open_ parameters (see Methods) highlighted that accuracy was mostly dependent on average promoter accessibility across cells and number of cells in the dataset (**Figures 2C and S3**). Most importantly, it underscored the very high accuracy of Enhlink (f1-score > 0.8) when enough cells were used (> 5000) or when a promoter was widely accessible (average accessibility > 0.2; **Figure 2C**).

**Figure 3.**
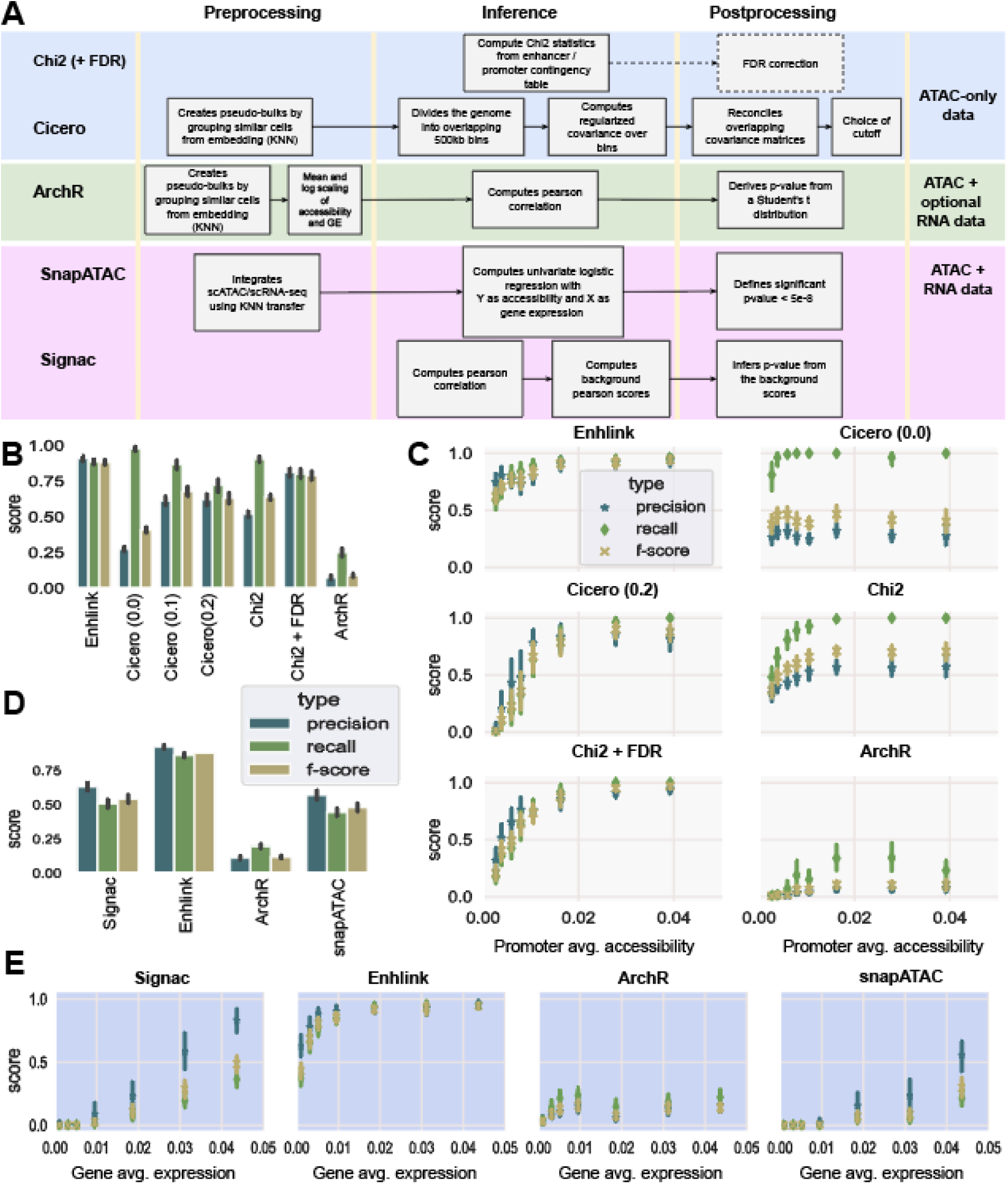
Enhlink outperforms other strategies for inferring linkage on simulated data. **A** Summary of existing enhancer-promoter method workflows. Some methods use scATAC-seq only as input (Cicero, Chi2 + FDR), others use scATAC-seq combined with scRNA-seq (Signac, SnapATAC). ArchR has a mechanism for both cases. **B** Enhlink outperforms ATAC-only methods on 400 simulated promoters and 1800 simulated enhancers generated from scATAC-seq data. The scores are computed from the average performance from each simulated promoter (see Methods). **C** Enhlink outperforms other ATAC-only methods independently of the promoter accessibility. Accuracy is dependent on the promoter accessibility (x axis) with more accessible promoters leading to better f1-scores. **D** Enhlink outperforms ATAC + RNA methods on 897 simulated genes and 4090 simulated enhancers inferred from the multiome snRNA-/snATAC-seq data. **E** Enhlink outperforms other ATAC + RNA methods across average gene expression values. Accuracy is dependent on the gene expression (x axis) with more expressed genes leading to better f1-scores (y axis).

### Enhlink outperforms other methods on simulated datasets

We compared Enhlink performance with that of other popular co-accessibility approaches implemented in Cicero (Pliner *et al*., 2018; Fang *et al*., 2021), SnapATAC (Fang *et al*., 2021), Signac (Stuart *et al*., 2021), and ArchR (Granja *et al*., 2021) (**Figure 3A**). We also performed a contingency table analysis relating a promoter and enhancer (Chi2) and, optionally, corrected the resulting *p*-value for genome-wide multiple hypothesis testing using the method of Benjamini and Hochberg (*Website*, no date a) (Chi2+FDR). Cicero and the Chi2 approaches are applicable to snATAC-seq data only, while SnapATAC (Fang *et al*., 2021) and Signac (Fang *et al*., 2021; Stuart *et al*., 2021) were applied to correlate enhancers with gene expression. ArchR was applied to both snATAC-seq data only and to correlate genes (continuous values) with enhancers (binary). We simulated one dataset of promoter and associated enhancer accessibilities, using the framework described above, and a second dataset of gene expression and associated enhancer accessibilities, using a similar framework (see Methods). Both datasets were derived from cell type (i.e., aCM or vCM)-specific λ_close_ and λ_open_ parameters and, thus, effectively simulate cells of a single cell type. We applied each method to the same set of query promoters and enhancers (see Methods). Enhlink outperformed other methods in terms of f1-score and precision computed for each gene/promoter (**Figures 3B-C**), with Chi2+FDR and Cicero (with a score cutoff of 0.2) matching Enhlink performance only for more accessible promoters (average accessibility > 0.025; **Figure 3D**). In the experiment with simulated gene expression, Enhlink performance (f1-scores ∼ 0.88) greatly exceeded those of Signac and SnapATAC (f1-scores ∼ 0.50) (**Figure 3E**). We considered modifications to ArchR, including changing parameters for the embedding step and replacing its *p*-value calculation with the one used by Signac. The results showed that using fewer neighbors (see Methods) increased the accuracy of ArchR with simulated genes (**Figure S4**), and suggested that the first embedding step of ArchR was actually detrimental to its accuracy.

### Enhlink enhancer-promoter associations are enriched for physical interactions

We next applied Enhlink to scATAC-seq and (promoter capture) PCHi-C data generated across two tissues – pancreatic islets and adipose – in our previous T2D study (Poirion *et al.,* 2023). This study examined the effect of mouse genotype (i.e., strain), sex, and diet on the cellular heterogeneity of these metabolic tissues in mice fed an obesogenic or laboratory diet. In both tissues, accessibility profiles clearly separated major cell types (**Figures 4A and 4C**), and we previously reported differences between genotypes and diets (Poirion *et al.,* 2023). Here, we applied Enhlink, Chi2 + FDR, and Cicero independently to each tissue and cell type to identify OCRs co-accessible with promoters. PCHi-C was previously performed on these same tissues to identify physical enhancer-promoter interactions (**see Availability of data and materials**). In both tissues, we observed higher recalls, precisions, and f1-scores (see Methods) for Enhlink compared to Cicero, and much higher numbers of links leading to higher recalls but lower precision for Chi2 + FDR compared to Enhlink (**Figures 4B and 4D**). Here, we did not apply a cutoff to filter Cicero links, so as to obtain a number of links similar to that resulting from Enhlink. Further, we found that Enhlink-inferred links were more likely to be shared across sequencing batches (as measured by entropy computed across batches, see Methods), and hence less likely to be induced by technical artifacts, than were those inferred by Cicero or Chi2 + FDR (**Figure 4F**). Indeed, the proportion of links with *zero* or close to zero entropy, indicating a batch-specific link and a possible batch effect, was lower amongst those inferred from Enhlink relative to the other two methods (**Figure S5**). Finally, we found that Enhlink-inferred links present in the PCHi-C data had lower *p*-values than those that did not, across all cell types (**Figure 4E**). This suggests that Enhlink *p*-values are a good indicator of the biological meaningfulness of a given link. As such, we generated an atlas of enhancer-promoter co-accessibilities for both tissues and each cell type (**Figure S6**), indicating genotype, sex, and diet-specific effects and annotated these links with PCHi-C interactions (**see Availability of data and materials**).

**Figure 4.**
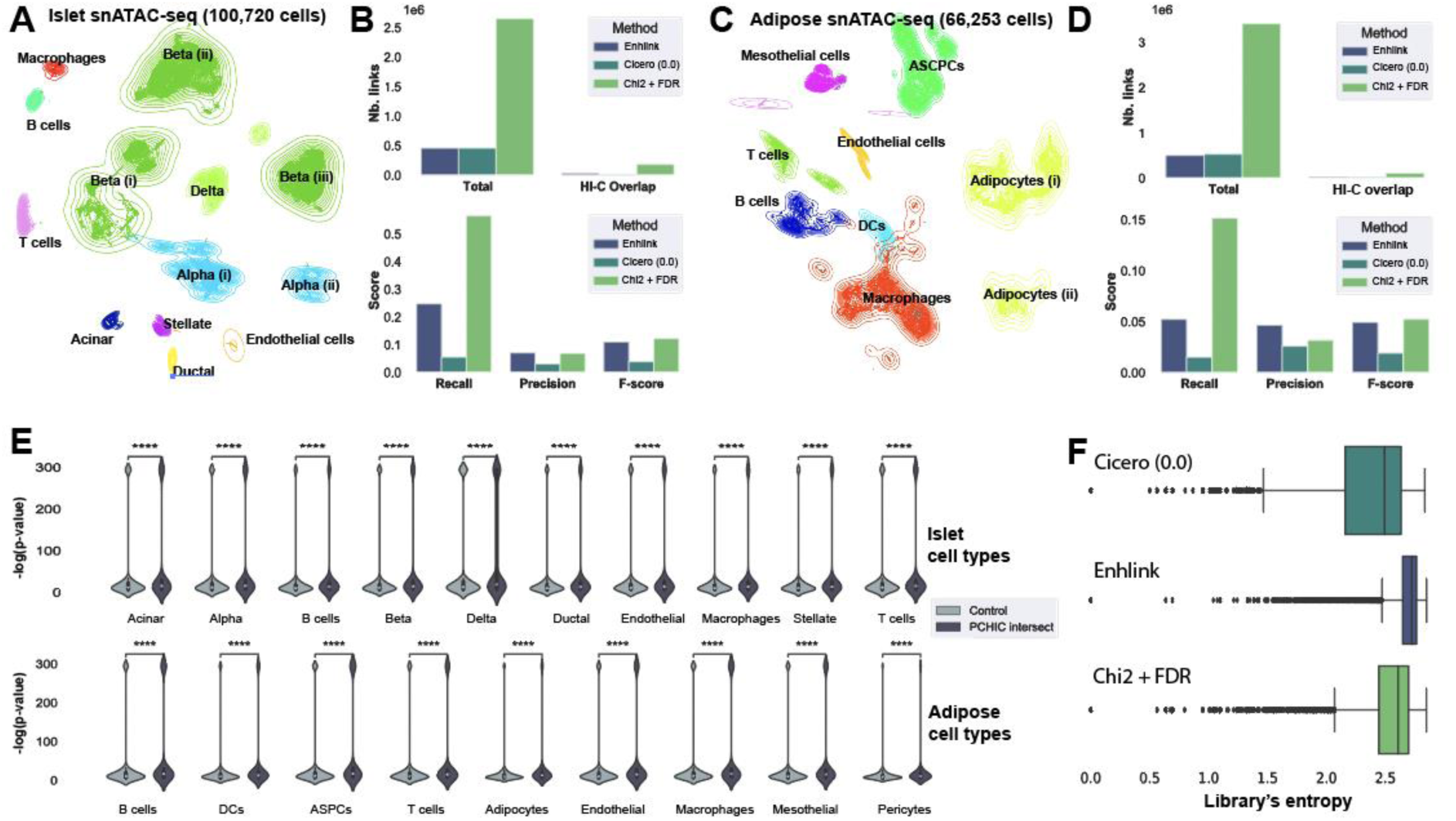
Enhlink outperforms other approaches in retrieving PCHi-C links and mitigates batch effects. **A** UMAP embedding and cell types of the islet dataset. **B** Enlink, Cicero, and Chi2 performance of promoter-enhancer inference in islet snATAC-seq relative to islet PCHi-C. **C** UMAP embedding and cell types of the adipose dataset. **D** Enlink, Cicero, and Chi2 performance of promoter-enhancer inference in adipose snATAC-seq relative to adipose PCHi-C. **E** Comparison (Mann-Whitney test) of the Enhlink *p*-value distributions from links intersecting PCHi-C and those not intersecting (control). **F** Distribution of the *batch x link* entropy for Cicero, Chi2 and Enhlink from a subset of cells from the islet dataset. Low entropy close to zero indicates links that exist only in a few or a single batch while high entropy indicates links widespread amongst the batches.

### Prioritizing neuronal enhancer-promoter links through multi-omic integration

To characterize epigenomic regulation within another heterogeneous tissue, we generated a multi-omic snATAC-/snRNA-seq dataset from the striatum region of the brain. Striata were collected from eight genetically diverse inbred mouse strains, representing the founders of the Diversity Outbred (DO) and Collaborative Cross populations used for genetic analysis of complex traits (Saul *et al*., 2019). Of particular interest in the striatum are a subset of striatonigral and striatopallidal neurons that are defined by expressing either the dopamine receptor 1 (*Drd1*) or 2 (*Drd2*), respectively (Gerfen *et al*., 1990). Clustering of gene expression identified eight major cell types and mixed populations of neurons, including the *Drd1*/*Drd2* neurons and all other previously identified major cell types (**Figure S7A).**

We sought to prioritize links between enhancers and promoters/genes based on both chromatin accessibility and gene expression data. Focusing first on *Drd1* neurons, we used Enhlink to identify 47,682 enhancer-promoter links within scATAC-seq data and 44,101 enhancer-gene links within scRNA-seq data. A subset of 16,431 links were concordant (i.e., shared and in the same direction), and these had higher scores and lower *p*-values than those uniquely identified from co-accessibility alone (snATAC-seq only; Mann-Withney *p*-value=0; Figures S7B-C). Similarly, enhancers within *Drd2* neurons with concordant (*n*=17,098) associations with a promoter’s accessibility (*n*=45,424) and its gene’s expression (*n*=47,023) had higher scores and lower *p*-values than those supported by co-accessibility alone (Figures S7B-C). Collectively, these results suggest that joint multi-omic analysis further refines enhancer identification.

We next used the expected association between a gene’s expression and its enhancers’ accessibility in the multi-omic data to evaluate computationally inferred co-accessibility between the gene’s promoter and those enhancers. For each marker gene of the Drd1 or Drd2 neurons, we inferred enhancer links with the associated promoter. We then evaluated the links according to whether they exhibited the expected correlation with the gene’s expression, as assessed with logistic regression (see Methods). Both Cicero (without cutoff) and Enhlink identified enhancer associations for 162 of the 172 marker genes. However, Enhlink identified a more focused set of links (*n*=802) relative to Cicero (*n*=6,997), having significantly stronger associations with gene expression (Mann-Whitney *p*-value<0.05) (Figure S8). This suggests Enhlink associations are enriched for true positives. Increasing Cicero’s cutoff resulted in smaller sets of links having increased correlation with gene expression relative to results without a cutoff, yet still inferior or similar to correlation inferred by Enhlink (Mann-Whitney *p*-value<0.05). Further, this came at the expense of *fewer* linked genes – 92 for the often used (Wang *et al*., 2020; Hocker *et al*., 2021) cutoff of 0.2 and 55 for the cutoff of 0.28 yielding the number of links (*n*=700) most similar to those obtained with Enhlink. Overall, these results show the strength of Enhlink in identifying putative enhancers more strongly associated with target gene expression relative to those identified by Cicero.

### Validating neuronal enhancer-promoter links with eQTL

One method to validate putative enhancers is through direct genome editing to test the downstream effect on gene expression. As an alternative approach, genetic variation among individuals provides a natural source of variation within enhancer sequences. The Diversity Outbred mouse population, segregates greater than 50 million variants with extremely high precision for genetic mapping (*Website*, no date b; Churchill *et al*., 2012; Saul *et al*., 2019). To explore the ability of Enhlink to identify biologically-relevant enhancers, we integrated previously collected (see Methods) expression quantitative trait loci (eQTL) data from bulk striatum RNA-seq experiments with enhancer links identified here. Genes with local (or *cis*-) acting eQTL are abundant in the DO population. Further, these *cis*-eQTLs are likely driven by variants within an enhancer proximal to the regulated gene that impacts expression (Wang *et al*., 2020), which we hypothesize would alter cell-type specific OCRs identified through snATAC-seq. Because the snATAC-seq data collected here represent replicates from the eight parental strains of the DO, we can further estimate OCR accessibility within each strain and see which accessibility patterns match the eQTL results. We identified 1,731 and 429 links from joint analysis of the snRNA- and snATAC-seq that were significantly (Enhlink *p*-value < 0.01) associated with Drd1 or Drd2 in both modalities, respectively (see Methods). To look for enhancers that drive differential expression between these two sub-classes of neurons we compared these links with the set of marker genes from Drd1/Drd2 neuron gene expression clusters (see Methods). Doing so identified 159 links for 66 genes for *Drd1* neurons and 32 links for 17 genes for *Drd2* neurons. Of the enhancers identified with Enhlink and associated with marker genes of *Drd1* or *Drd2* neurons, 68 are linked to genes with *cis*-acting eQTL. We performed a SNP association analysis using a logarithm of the odds (LOD) regression approach to identify variants that show an association between the genotype at the OCR and gene expression (see Methods). Candidate enhancers driving variation in expression were identified as those with matching correlations between their genotype at the variant, gene expression, and accessibility of the promoter and enhancers across inbred strains. Of the identified correlations, three enhancer-promoter links for marker genes *Gulp1*, *Kcnb2*, and *Col25a1* (**Figures 5A-B**) serve as proof of principle. In each case the strains with the genotype identified as having the largest effect on the eQTL from bulk data (alternative genotype, **Figure 5C**), presented differential accessibility and expression in the single-nuclei data compared to the other strains (**Figure 5B**). Together these data show that Enhlink identifies biologically relevant enhancers that play an active role in cell-type- and strain-specific gene regulation.

**Figure 5.**
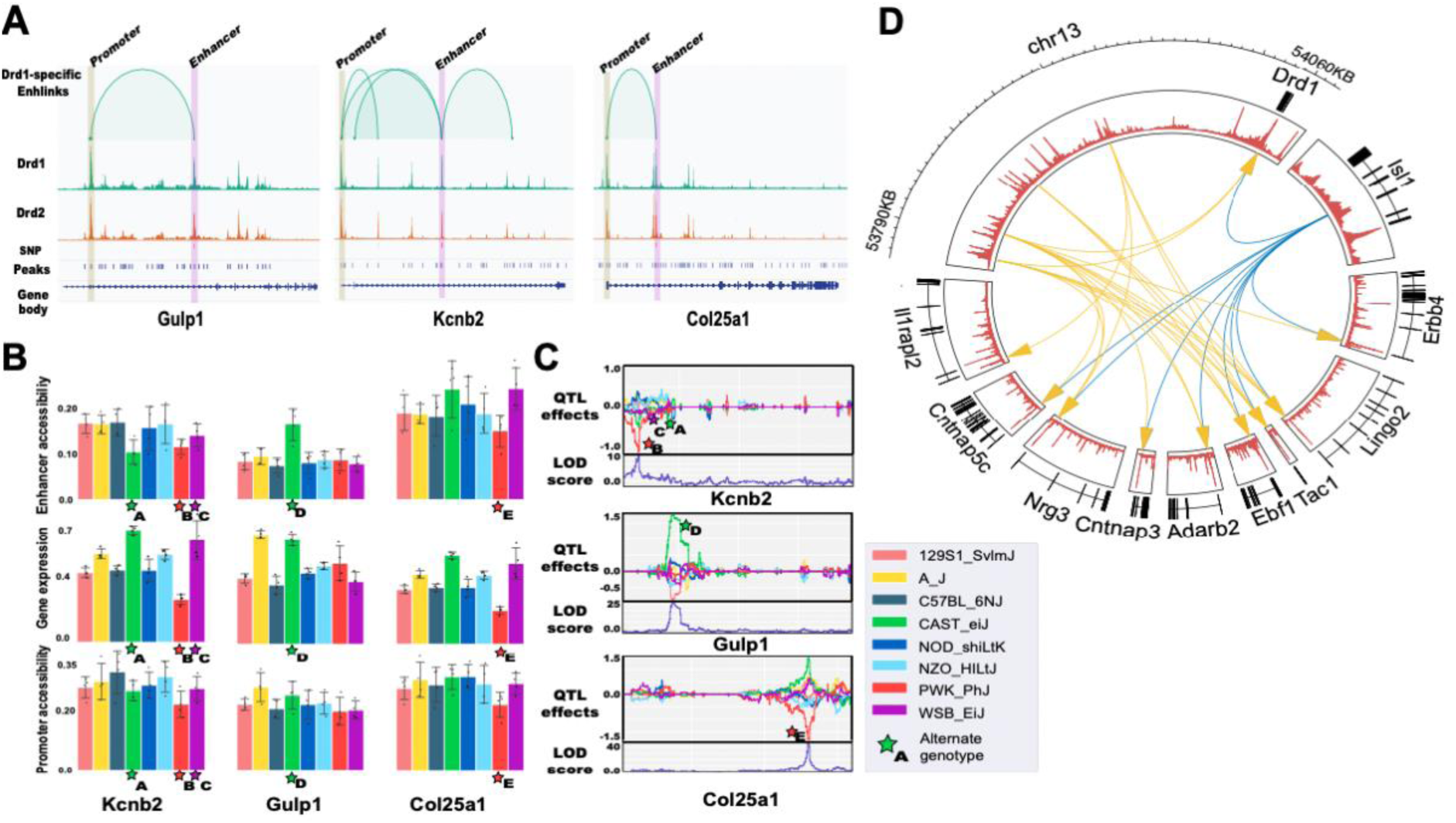
Enhlink reveals chromatin regulation mechanisms of striatum Drd1/Drd2 neurons. **A** Chromatin accessibility (y axis) with Enhlink-inferred links between the promoters and enhancers for *Kcnb2*, *Gulp1*, and *Col25a1*, three marker genes of Drd1 neurons. **B** Chromatin accessibility and gene expression profiles per genotype for three enhancers (*Kcnb2*, *Gulp1*, and *Col25a1*). **C** eQTL logarithm of odds (LOD) scores for SNPs within the boundaries of the three enhancers across the eight DO genotypes. Stars indicate genotype harboring an alternative allele within an enhancer of *Kcnb2*, *Gulp1*, or *Col25a1*. Star subscript associates LOD scores in panel C with chromatin accessibility and gene expression in panel B. **D** Distal Enhlink analysis unveils multiple enhancers from the region 500kb downstream of the *Drd1* promoter and linked to the top 10 marker genes of *Drd1* neurons (yellow arrows). These genes are also linked to an intronic region of *Isl1* (blue arrows), a key gene regulating Drd1/Drd2 genetic programs.

### Identifying distal *Drd1*-specific enhancers

While the above analysis focused on identifying how local genetic variation impacts gene expression, Enhlink can also be used to identify cell-type-specific *distal* enhancer networks. To do so we extended our analysis by finding distal common enhancers for the top 10 *Drd1*-specific upregulated marker genes (*Adarb2, Cntnap3, Lingo2, Drd1, Il1rapl2, Cntnap5c, Erbb4, Nrg3, Tac1, Ebf1*) and top 10 *Drd2* specific marker genes (*Drd2, Nell1, Necab1, Unc5d, Grik3, Ptprm, Fam155a, Chrm3, Penk, Adk*) across *all* (259,720) OCRs, rather than those within +/-250kb of their respective promoters. We identified 110 links from snATAC-seq data alone. To mitigate potential false positives over this genome-wide set of OCRs, we further applied Enhlink to the multi-omic snATAC-/snRNA-seq data as above and inferred 70 links. We prioritized the 33 links shared across both analyses for further analysis. Remarkably, 22 of these 33 links were associated with just four enhancers, all arising from a region 500 kb downstream of the *Drd1* promoter and surrounded by the predicted pseudo-genes: *Gm34439*, *Gm40954*, and *Gm34557* (**Figures 5D** and **S9A**). Further, all 10 of the *Drd1*-specific genes have at least a distal enhancer within the four *Drd1* proximal enhancers (**Figures 5D** and **S9A**). Additionally, nine out of the remaining 11 links matched an intergenic OCR from the Islet-1 (*Isl1*) gene that was previously characterized for regulating striatonigral and striatopallidal genetic programs (Lu *et al*., 2014) (**Figures 5E** and **S9B**). Notably, neurons from an *Isl1* knockout mouse showed an increase in *Drd2* expression and promoted striatopallidal (*Drd2*) neuron differentiation while repressing striatopallidal (*Drd1*) genes. Together these data show that Enhlink can identify coordinated chromatin regulation at distal loci with biologically meaningful connections, perhaps indicating coordinated transcription factor activity (Yang *et al*., 2017).

## Discussion

In this study, we introduce Enhlink, a novel computational method that efficiently infers genomic linkages from single-cell datasets and is suitable for complex experimental designs. Enhlink infers enhancer-promoter co-accessibility from chromatin accessibility profiles in scATAC-seq data. It can also infer enhancer-promoter links supported by both their co-accessibility and concordant enhancer accessibility and target gene expression within multi-omic snATAC-/snRNA-seq datasets. More generally, Enhlink could be applied to other single-cell modalities containing sparse high-dimensional data, such as single-cell DNA methylation (Liu *et al*., 2021), single-cell ChIP-seq (Grosselin *et al*., 2019), or multi-omic datasets combining epigenome, methylome, and/or transcriptome (Dimitriu *et al*., 2022).

Enhlink leverages an original procedure derived from random forests (Breiman, 2001) to extend the capabilities of existing methods. First, Enhlink adjusts for technical covariates, such as sequencing library ID, to minimize batch effects at the level of *individual* cells. Methods that instead apply batch correction to aggregates of similar cells do not readily accommodate technical effects *within* the aggregate. This would pose a problem in studies, such as our T2D study, where cells of similar genotype are partitioned across batches (see Methods). Second, Enhlink can infer enhancers linked to a promoter within specific contexts, such as sex, genotype, or disease, by including each as a biological covariate. Third, for each enhancer-promoter link, Enhlink infers a *p*-value that is adjusted for multiple hypothesis testing *relative to all and only those enhancers tested for association with that promoter*. In this way, the strength (i.e., adjusted *p*-value) of each inferred association is scaled according to the genomic context of each promoter. Hence, two enhancers may have different inferred associations, even if their correlations with their respective promoters are similar, for example, if one enhancer has a much higher correlation with other enhancers it is compared against than the other enhancer. Fourth, Enhlink can infer linkages from distal regions beyond the neighboring OCRs. Finally, Enhlink can perform power analyses to estimate expected accuracies based on a simulation workflow developed from experimentally validated enhancer-promoter linkages. This simulation framework can be applied independently of Enhlink and can aid others in diagnosing the impact of hyperparameter tuning, preprocessing steps, or other methodological choices.

Enhlink employs several regularization mechanisms to reduce false positives. One hyperparameter, the maximum number of explanatory features of each tree, is biologically interpretable as the *expected* number of enhancers at each target region. While the actual number of enhancers is a property of the entire *ensemble* of trees, we expect that it is of the same order of magnitude as the number of features considered at each *tree*. As such, the hyperparameter can be set according to biological expectations or tractability of downstream experimental validation. Enhlink sensitivity and specificity can also be fine-tuned by adjusting the number of trees used and the minimum number of features required for each tree. Thus, weak but significant enhancer-promoter associations could be inferred by using a larger forest and feature size at the expense of increased computation. We implemented Enhlink in Go which allows it to be extensively distributed amongst a cluster of CPUs and makes it well-suited for analyzing current (Zhang *et al*., 2021) and future (Vandereyken *et al*., 2023) large-scale multi-omic single-cell datasets.

Through extensive benchmarking of both simulated and real data, our study demonstrates that Enhlink significantly outperforms other existing approaches in terms of accuracy while also limiting artifactual, batch-specific linkages. We took advantage of multi-omic data to show that enhancers identified from scATAC-seq were better correlated with gene expression when inferred with Enhlink than with the popular Cicero framework. In addition, simulation and intersection with two reference PCHi-C datasets showed that Cicero was also outperformed by a Chi2 procedure followed by FDR correction. We also showed that aggregating cells into pseudo-bulk, a preprocessing step followed by ArchR and inspired by the Cicero workflow, was actually detrimental to accuracy. This occurs when the binning, aimed to group cells with similar characteristics, is too coarse and results in the grouping of heterogeneous cells. We evaluated Enhlink using three datasets, including two snATAC-seq datasets previously generated from mouse islet and adipose tissues and a novel multi-omic snATAC-/snRNA-seq dataset from the mouse striatum. The single-cell islet and adipose studies aimed to characterize cellular heterogeneity in these tissues and uncover genes and regulatory elements perturbed by diet and genetic mechanisms. To aid this investigation, we developed cell-type-specific enhancer-promoter atlases of the islet and adipose tissues (see **Availability of data and materials**). These include genotype-, diet-, and sex-specific linkages that are annotated according to whether the corresponding promoter and enhancer physically interact based on PCHi-C data generated from the same tissues. Links supported by PCHi-C data, and those with concordant promoter chromatin accessibility and target gene expression in the striatum dataset, have significantly lower Enhlink *p*-values than their counterparts. This suggests that Enhlink *p*-values robustly reflect the biological meaningfulness of their associated links.

We generated the multi-omic mouse striatum dataset to investigate epigenomic regulations underlying the differentiation and gene regulation of striatonigral neurons expressing the dopamine receptor *Drd1* and striatopallidal neurons expressing *Drd2*. Our goal was to identify strong candidate regulatory regions involved in these processes. To achieve this, we utilized Enhlink to identify concordant links between promoter accessibility and gene expression. We then examined the relationship between naturally occurring regulatory variation, chromatin accessibility, and gene expression amongst eight parental haplotypes with associated eQTL and SNP data. Through this analysis, we identified three enhancers associated with the expression of *Gulp1*, *Col25a1*, and *Kcnb2*. The contribution of the different haplotypes to the expression pattern indicated that the variants located within the Enhlink-identified enhancers were likely to be the causal factors for the observed changes in accessibility and expression. This finding strengthens the hypothesis that the regions identified by Enhlink play a crucial role as enhancers. Given that the majority of disease-associated variants within the human population occur within OCRs, this approach is extendable to prioritizing variants related to complex traits and might palliate some of the theoretical limits of eQTL analysis (Umans, Battle and Gilad, 2021).

We further demonstrated the power of Enhlink to detect links between a target promoter and its distal enhancers (> +/-250kb). We hypothesized that key enhancers should be detected as hubs correlating with the expression of multiple genes, similar to other studies modelling gene expression with SNPs (Gamazon *et al*., 2015; Poirion *et al*., 2018). Strikingly, we found that most of the distal links inferred from the top marker genes of the striatonigral neurons came from a region located 500kb downstream of the *Drd1* promoter. This region exhibits many characteristics of a super-enhancer region (Heinz *et al*., 2015), as it contains a cluster of enhancers associated with several marker genes of the striatonigral neurons. Super-enhancers have been shown to determine cell fate and to maintain cell identity (*Website*, no date c), stressing the possible role played by the downstream region of *Drd1*. Also, we found that all top ten markers of *Drd1* neurons were linked with an intronic region of the *Isl1* gene, a key gene in the striatopallidal/striatonigral differentiation (Lu *et al*., 2014). While these results are novel, they need to be confirmed through further experimental validation.

Enhlink is a powerful and robust method for inferring genomic linkages from sparse, high-dimensional, single-cell ‘omic or multi-omic datasets. Enhlink outperforms other tested approaches, in part, by introducing cellular level covariates that ameliorate technical effects and capture biological effects. Enhlink’s efficient implementation is tailored to large-scale single-cell analyses, including those aimed at deciphering complex regulatory networks.

## Methods

### Human heart snATAC-seq dataset processing

We downloaded the Human Heart snATAC-seq dataset from the portal (http://ns104190.ip-147-135-44.us/CARE_portal/ATAC_data_and_download.html) described in the publication (Hocker *et al*., 2021). From the portal, we downloaded the matrix file (*all.npz*), the cell index file (*all.index*), the OCR features index file (*all.merged.ygi*), the genome reference (Homo_sapiens.GRCh38.99.TSS.2K.bed) the cluster file (*all.cluster*), and the *all.group* file with the library ID of each cell. The matrix from the *all.npz* file is a scipy sparse matrix with *79515* cells and *287415* OCRs with boolean values indicating if at least one read mapped to the cell is found within the boundaries of the corresponding OCR. From the genome reference and features index files, we defined the *KCNH2* promoter regions by the following OCRs: “chr7:150976584-150977120”, “chr7:150978193-150978805”, and “chr7:150978915-150979661”. The *KCNH2* enhancer was defined as the following genomic region: “chr7:150955147-150956502”. We also defined the *MYL2* promoter regions by “chr12:110920282-110920944” and “chr12:110921386-110921633”. We defined three enhancer regions of *MYL2* by “chr12:110931149-110931877”, “chr12:110928658-110929096”, and “chr12:110907461-110909456”. We then focused on the atrial (aCM) and ventricular (vCM) cardiomyocyte cell types, since *KCNH2* is only expressed in these cell types, and defined a *KCNH2* promoter boolean vector for either aCM or vCM by merging (using a *logical or* operand) the three promoter region vectors from the feature matrix using either the cell index of aCM or vCM. We followed the same strategy for the promoter of *MYL2*, but restricted to the vCM cell type since *MYL2* is only expressed in vCM. We also downloaded the bigwig tracks from the same portal for the aCM and vCM cell types from the same portal. Finally, we computed the Enhlink co-accessibilities for these two promoters and for aCM and vCM based on the workflow described below.

### Co-accessibility signals from *KCNH2* and *MYL2*

We used the *KCNH2* promoter vectors of aCM and vCM as ground truth labels (either accessible or not for a given cell) and the *KCNH2* enhancer vectors as estimated labels to compute the precision, recall, and f1-scores. We then drew a random subset of 500 enhancers amongst the 287415 features of the feature matrix to compute the baseline distributions for the precision, recall, and f1-score. We then derived a *p*-value for the computed precision, recall, and f1-score with regard to the baseline distributions. We followed the same strategy for the *MYL2* promoter and its associated enhancers.

### Mouse islet and adipose scATAC-seq

We used two scATAC-seq datasets from the mouse islet and adipose tissues whose generation and processing we previously described (Poirion *et al.,* 2023). The islet dataset gathered 100,720 cells, 295,089 OCR features, ten cell populations including alpha, beta, and delta cells, and was processed using 18 10X Genomics sequencing libraries. The adipose dataset gathered 60,229 cells, 311,645 OCR features, nine cell populations including adipocytes and macrophages, and was processed using 24 10X Genomics sequencing libraries. Cells from both datasets were labeled with their mouse strain/genotype ID (CAST, NZO, B6), diet (10% or 44% fat), and sex (M/F).

### Mouse (RNA/ATAC) multi-omic single-cell collection

Mouse striatum were dissected from 12-week-old mice from eight inbred strains: A/J (The Jackson Laboratory Stock 000646), C57BL/6J (000664), 129S1/SvImJ (002448), NOD/ShiLtJ (001976), NZO/H1LtJ (002105), CAST/EiJ (000928), PWK/PhJ (003715), and WSB/EiJ (001145). Striatum were collected from two males and two females for each strain. Striatum samples were flash frozen in liquid nitrogen and stored at -80 °C until processing. After collection, the striatum samples were processed in four batches of eight over two days. Each batch consisted of four male and four female samples, and each strain was represented in each batch.

### Multi-omic library generation

For single nuclei preparation, frozen striatum were placed into 500 µl Miltenyi Nuclei Extraction Buffer (Miltenyi 130-128-024) plus 0.8 U/µl RiboLock RNase Inhibitor (ThermoFisher EO0382) in a gentleMACS C tube (Miltenyi 130-093-237) on ice, and then nuclei were immediately extracted in batches of 4 through the “4C_nuclei_1” program on the MACS Dissociator. Nuclei were placed on ice and filtered through a 70 µm pluriStrainer Mini into a pre-chilled 2 ml tube, and the C tube was washed with an additional 500 µl cold Miltenyi Nuclei Extraction Buffer and filtered. Nuclei were spun down at 350 rcf at 4 °C for 5 minutes and resuspended in 80 µl cold PBS with 2% BSA and 1 U/µl RiboLock. Nuclei were counted (∼4,000 nuclei/µl) and 60,000 nuclei from each of the 8 samples were mixed in a chilled 1.5 ml Eppendorf tube. The pooled nuclei were spun at 300 rcf at 4 °C for 3 minutes and resuspended in 30 µl 10x Genomics Nuclei Buffer with 1 U/µl RNase inhibitor. The pooled nuclei were counted once more to confirm counts (∼10,000 nuclei/ul), single-cell suspension, and lack of debris.

Nuclei viability was assessed on a LUNA-FX7 automated cell counter (Logos Biosystems), and up to 40,000 nuclei (∼5,000 from each sample) were loaded onto each of 4 lanes of a 10X Chromium microfluidic chip. Single nuclei capture and library preparation were performed using the 10X Chromium platform and according to the manufacturer’s protocol (#CG000388 Chromium Next GEM Single Cell Multiome ATAC + Gene Expression). Because the 10X chip was superloaded, to reduce duplicate reads arising from multiple PCR steps, the number of cycles was adjusted from the 10X protocol. After GEM generation, barcoded cDNA and transposed DNA fragments were pre-amplified using 6 cycles. The pre-amplified sample was divided and used for two separate steps. An additional 6 cycles were used for adding a sample index for ATAC library construction and 12 PCR cycles for the gene expression library construction. cDNA and ATAC libraries were checked for quality on Agilent 4200 Tapestation and ThermoFisher Qubit Fluorometer, and quantified by KAPA qPCR, before sequencing; each gene expression library was sequenced using NovaSeq 6000 S4 v1.5 200 cycle flow cell lane, dual index scRNAseq asymmetric read configuration 28-10-10-90, targeting 20,000 nuclei with an average sequencing depth of 50,000 read pairs per nucleus. Each ATAC library was sequenced using Illumina NovaSeq 6000 S2 v1.5 100 cycle flow cell lane, with a 50 -8-24-49 read configuration, also targeting 20,000 nuclei with an average sequencing depth of 50,000 reads per cell.

### Multi-omic library sequencing

Illumina base call files for all libraries were demultiplexed and converted to FASTQ files using bcl2fastq 2.20.0.422 (Illumina). A filtered joint digital gene expression and chromatin accessibility matrix was generated against the 10x Genomics mm10-2020-A reference build (version 2020-A, Assembly: GRCm38, ENSEMBL release 98; Annotations: Gencode vM23) using a modified 10x Genomics CellRanger-ARC count pipeline (v2.0.0), which had the 20,000 cell limit of cell calling removed.

### scRNA-seq analysis for the striatum dataset

For each batch of the libraries, the output from Cell Ranger ARC consisted of both ATAC and gene expression BAM files. These two BAM files were used as inputs for Demuxlet (Kang *et al*., 2020) to determine the mouse strain identities of individual cells and to detect doublets. Only cells that were identified as singlets with consistent mouse strain identities in both assays were retained for further analysis.

To annotate and visualize the retained single cells, we performed downstream gene expression analysis on them using Seurat (version 4.0.3, (Hao *et al*., 2021)). For each batch, we removed low-quality cells and multiplets by filtering out cells with fewer than 200 or more than 7,500 detected genes and those with greater than 15% mitochondrial counts. We then used DoubletFinder (McGinnis, Murrow and Gartner, 2019) to further exclude any remaining doublets. Next, we merged the Seurat objects from the four batches and performed normalization, highly variable feature identification, scaling, and linear dimensionality reduction using the NormalizeData(), FindVariableFeatures(), ScaleData(), and RunPCA() commands, respectively, with default parameters. To remove batch effects, we integrated the single-cell datasets from the four batches using Harmony (Korsunsky *et al*., 2019). Using the first 40 principal components determined manually by the Elbow plot, we conducted unsupervised cell clustering through the Louvain algorithm on the K-nearest neighbor (KNN) graph with resolution set to 0.2, which resulted in 22 cell clusters. We visualized the output in a 2D Uniform Manifold Approximation and Projection (UMAP) embedding using the same PCs used for cell clustering. We excluded four cell clusters that (1) co-expressed neuron markers *Drd1/Drd2* and oligodendrocyte marker *Aspa*; (2) co-expressed neuron markers *Drd1/Drd2* and astrocyte marker *Gja1*; (3) co-expressed neuron markers *Drd1/Drd2* and macrophage marker *C1qa*; or (4) highly expressed mitochondrial genes. We then re-processed and integrated the remaining cells using the same steps as before, yielding 18 cell clusters. We annotated these 18 cell clusters using marker genes identified through the FindAllMarkers() command through DropViz (Saunders *et al*., 2018).

We processed the snATAC-seq data separately from the scRNA-seq data and followed the same preprocessing procedure as the one described for the mouse islet and adipose snATAC-seq dataset (Poirion *et al.,* 2023). Briefly, we aligned the reads to the mm10 genome with Cell Ranger V6 with default parameters. We inferred the peaks fusing MAC2 (Zhang *et al*., 2008).

### Enhlink analytical workflow

The analytical procedure carried out by Enhlink involves multiple steps illustrated in **Figure S1** and explained in greater detail in Additional File S1. It can be summarized as follows: a) create a feature matrix (i.e., OCR x cell matrix) and a response vector (i.e., single-cell promoter accessibility or target gene expression) for each target genomic region, b) model the response vector as a function of the feature matrix and identify the significant features associated with the target region, and c) optionally perform a secondary analysis to detect biological covariates associated with the linkage.

At a minimum, Enhlink requires a boolean sparse matrix, indicating the OCRs of each cell and a list of genomic regions, typically the promoters, defining the linkage targets. In addition, Enhlink optionally takes a sparse matrix containing the values of the target regions (such as a single-cell expression matrix), a file containing the covariates of each cell, and the cluster IDs of each cell. Enhlink constructs a feature matrix *M*_*n*_ for each target region by iterating through the OCRs surrounding it (+/- 250kb by default). If the feature matrix contains fewer than 100 features by default, Enhlink supplements it with random features derived from the existing features. After constructing the feature matrix for each target region, Enhlink incorporates one-hot-encoded covariates and generates a response vector *v*_*g*_, representing either the boolean accessibility or the expression of the gene/target region *g*. If *v*_*g*_ is continuous (e.g., representing gene expression), it is binarized using the mean of the non-null values as threshold. Then, Enhlink proceeds to model *v*_*g*_ = *f*(*M*_*n*_) for each cluster (or for all cells if no clusters are provided) and identifies features from Mn that significantly predict Vg. Enhlink employs a strategy similar to that of a random forest classifier (Breiman, 2001), performing 100 iterations by default. In each iteration, a random sample of cells and features is selected, and a decision tree (Quinlan, 1986) is used to recursively reduce the number of samples and features based on the top feature selected at each level of the tree. The top features are selected using a modified score derived from the Information Gain (IG) (see **Additional File S1**), which favors informative, positively correlated, and accessible features. For each feature, Enhlink calculates a *p*-value for its score to be different from zero across iterations using Student’s *t*-test from the disturb library (https://pkg.go.dev/gonum.org/v1/gonum/stat/distuv) and corrects the feature *p*-values using the Benjamini-Hochberg False Discovery Rate (FDR) procedure.

### Estimating the expected accuracy of inferred links through simulation

Enhlink provides an option to estimate the expected accuracy of the inferred links for a given target region by generating simulated enhancers and target regions. This simulation procedure aims to replicate the observed correlations between experimentally validated enhancers and promoters. More detailed information on this procedure can be found in Additional File S1. Briefly, Enhlink first simulates a promoter by shuffling a target region. Then, to simulate an associated enhancer, Enhlink duplicates the simulated promoter, and introduces two types of random noise. The noise is controlled by two hyperparameters, λ_open_ and λ_close_, which model the scenario where the enhancer is not accessible in a given cell (λ_open_) or the target region is not accessible (λ_close_). The values of λ_open_ and λ_close_ are estimated from the heart snATAC data, using the experimentally validated enhancer of *KCNH2* and enhancers from *MYL2*, as described below.

### Inferring biological context-specific enhancer-promoter interactions

Enhlink can optionally infer biological context-specific linkages with the details of the procedure found in Additional File S1 and summarized here. Each context is represented by a cell-level categorical covariate. Note that Enhlink one-hot encodes categorical variables into boolean features and cannot currently process continuous covariates. In this configuration. Briefly, Enhlink first infers the set of enhancers and covariates linked to a target region *g* then, for each of these enhancers *e* Enhlink computes *Veg* = *Ve* ∘ *Vg* as the Hadamard product between the target vector *Vg* and the enhancer vector *Ve. Veg* corresponds to a boolean vector indicating when both the enhancer *e* and the target region *g* are accessible within a cell. *Veg* is then used as a new target vector to find the covariates significantly associated with it.

### Addressing class imbalance in datasets with unequal covariate distributions

Enhlink provides an option to mitigate class imbalance in datasets with varying covariate distributions, which is beneficial when one or more covariates are under- or over-represented. In such cases, the bootstrap samples may not adequately represent the covariate space. Enhlink addresses this issue by generating a more balanced distribution of covariates through an incremental process. Specifically, Enhlink iteratively selects subsets of cells according to their covariates to obtain a near-uniform distribution of the covariates. While this strategy can be helpful in achieving a more uniform distribution of covariates, it is important to note that it may produce biased results if one or more covariates are vastly underrepresented. Therefore, it is recommended to thoroughly assess the distribution of covariates, and to consider alternative approaches such as covariate removal if necessary to ensure more representative results. More details are given in Additional File S1.

### Enhlink hyperparameters

The Enhlink inference workflow is governed by multiple hyperparameters summarized in **Table S1**. The main hyperparameters governing the regularization are *max_features*, which controls the maximum number of features that a tree can use, and *depth* which controls the maximum depth of each tree. *Depth* and *max_features* are intertwined since *depth* also indirectly controls the maximum number of features. However, *max_features* is a more intuitive hyperparameter to use than *depth*. The latter influences the speed of Enhlink and is set to 2 by default, but needs to be increased if *max_features* is set higher. *secondOrderMaxFeat* (set as 2 by default) is the equivalent of the *max_features* parameter for the inference of the biological context-specific enhancer-promoter interactions, triggered with the *secondOrder* hyperparameter. *N_boot* defines the number of trees to build and controls the false positive rate and influences the speed of Enhlink, similarly to *min_matsize*, controlling the minimum number of features of *Mn*. The hyperparameters *downsample* (the size of the bootstrap) and *maxFeatType* (the number of features to include) can be used to increase the speed of the procedure but lower values might lead to lower accuracies. Finally, *threshold* defines the *p*-value cutoff and can be used to regulate the precision and the recall.

### Enhlink software suite

Enhlink is an analytical framework developed in Go (https://go.dev/), and compiled into three executables: *enhlink, enhgrid,* and *enhtools*. The command line manual and arguments of each executable can be accessed using the *-h* flag, (e.g. *enhlink -h*). *enhlink* is the main executable that launches the Enhlink pipeline, while *enhgrid* allows launching Enhlink for a range of input values for all the hyperparameters accepting a numerical value. *enhgrid* is useful for, for example, automatizing a grid-search approach by trying a combination of multiple hyperparameters or for testing different noise levels. *enhtools* intersect results from multiple runs and output either the common or unique links of a particular run. It also computes the accuracy between two runs (f1-score, precision, recall). Finally, *enhtools* can filter links that are not within specific regions defined in an input BED file. This functionality can be used for example to filter links not inside topologically associated domains (TAD). The Go source code, the manual, and the tutorial are available here: (https://gitlab.com/Grouumf/enhlinktools).

### Simulation studies of the performance impact of hyperparameters

We conducted a simulation study to estimate the impact of the different hyperparameters using three scATAC-seq datasets: the mouse islet, the mouse adipose, and the snATAC-seq from the multi-omic striatum dataset mentioned earlier. We aimed to investigate the impact of various hyperparameters and experimental conditions, such as dataset origin and read depth, on Enhlink’s expected accuracy (see **Figure 2C**). To accomplish this, we first subset the datasets by selecting specific cell types, such as beta cells for the islet, adipocytes for the adipose, and Drd1 neurons for the striatum. We then utilized Enhlink’s simulation framework, described in **Additional file S1**, to estimate the expected accuracy of Enhlink analysis at a given promoter. We used a consistent grid of hyperparameters, as detailed in **Table S2**, for each dataset and applied the enhgrid tool to perform Enhlink on the hyperparameter grid for all three datasets. Furthermore, we conducted the Enhlink analysis on a random subset of 40 genes from the features list of the three datasets for each combination of hyperparameters in the grid.

### Estimating the hyperparameters importance from the simulation studies

From the hyperparameters simulation study described above, we estimated the most significant hyperparameters by fitting a Random Forest regressor from the scikit-learn library (Pedregosa *et al*., 2011) using the f1-score as the dependant variable, and Enhlink’s hyperparameters as predictive variables (**Figure S3**). The simulation generated the true positive rate (TPR), false positive rate (FPR) and false negative rate (FNR) for each gene tested and for each hyperparameters combination. We then computed the f1-score as 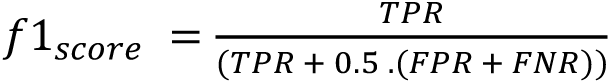 and model it as a function of 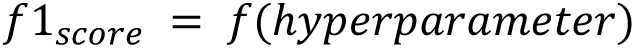. We one-hot encoded all the hyperparameters and their values, removing the first value to avoid colinearity, and used them as binary variables. We also added the dataset ID as an additional variable. We then collected the feature importance of the model, computed as the Gini importance, an impurity-based measurement, using the *feature_importances_* attribute of the scikit-learn class. For each simulated promoter, we also computed the accessibility ratio as the ratio between the number of cells having the promoter accessible and the total number of cells. We then plotted the influence of the promoter accessibility ratio on the f1-score using the Python library *seaborn*.

### Generating reference datasets for methods comparison

We generated two simulated datasets, using either snATAC-seq only from the mouse islet or the snATAC-seq + scRNA-seq from the striatum dataset, containing either simulated enhancer-promoter co-accessibilities (ATAC-seq only) or correlated enhancer-gene links (ATAC+RNA). From the mouse islet dataset, we used the delta cell subset and a random set of 400 genes to simulate 400 promoters and 1800 enhancers. For each random promoter, we generated a random number of simulated enhancers (between 2 and 7) with the noise parameters λ_close_*=1.25,* λ_open_*=0.25.* We created dummy genomic coordinates for these enhancers in order for them to be in the vicinity of their matching promoter. These simulated enhancers/promoters were injected in a matrix of 10100 cells (the delta cells) and 295089 peaks (the total number of peaks). From the striatum dataset, we used the Drd1 neuron cell types and a random set of 897 genes to simulate between 2 and 7 enhancers for each gene, for a total of 4090 enhancers. We used the same noise parameters: λ_close_*=1.25,* λ_open_*=0.25*, and binarised the randomized gene vectors using the mean of its non-null element before generating their associated enhancers (see above). The simulated enhancers were further injected into a matrix of 10000 cells and 259720 peaks.

### Methods comparison procedure

We processed the ATAC simulated matrix with Enhlink, Chi2, Chi2 + FDR, Cicero, and ArchR workflows, and processed the ATAC+RNA simulated matrix with Enhlink, Signac, SnapATAC, and ArchR workflows. We used the same genomic window of +/- 250kb around the promoter for all workflows. We then computed the 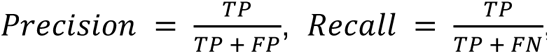, and the 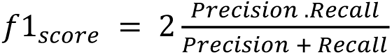 for each of the simulated promoters/genes using the set of simulated enhancers as the true positives. We computed for each workflow and dataset the overall accuracy scores with their standard deviation (sd) using the mean and the sd of the three metrics. Finally, we also plotted the correlation between the accuracy scores and the mean of each simulated gene or promoter using the *lmplot* function of the Python *seaborn* library.

### Alternative workflows implementation

#### Chi2

A straightforward approach for inferring the co-accessibility of a promoter region *g* with the set *E* of its surrounding enhancers is to perform a Chi2 test between *g* and *e* for each *e* ∈ *E*. Since *v*_*g*_ and *v*_*e*_, i.e. the accessibility vector of *g* and *e,* were binary, we constructed a *2 x 2* contingency table for each (*g, e),* containing the occurrence for the following conditions: *v*_*g*_ = = 1 ∩ *v*_*e*_ == 1, *v*_*g*_ == 0 ∩ *v*_*e*_ == 1, *v*_*g*_ == 0 ∩ *v*_*e*_ == 0, and *v*_*g*_ == 1 ∩ *v*_*e*_ == 0. We used the *chi2_contingency* function from the Python library *Scipy* (Seabold and Perktold, 2010) to infer the *p*-value.

#### Chi2 + FDR

A more refined approach to the Chi2 method described above is to correct for multiple hypothesis testing by applying the False Discovery Rate procedure from Benjamini-Hochberg (*Website*, no date a) on the set of *p*-values obtained when applying the Chi2 on *E.* The method consisted in computing an expected *p*-value (fdr), assuming a false positive, based on the rank of the feature and the corrected *p*-value was equal to *p-value x fdr*. We used the *fdrcorrection* method from the Python *Statsmodels* library (Seabold and Perktold, 2010) to perform the FDR corrections.

#### Cicero

We downloaded the latest version of the Cicero algorithm (Pliner *et al*., 2018) using the *R* library developed by the authors (https://cole-trapnell-lab.github.io/cicero-release/docs/). To parallelize the workflow and reduce the shared memory used, we processed each chromosome independently. For each chromosome, we created a UMAP embedding with the following steps: i) We performed a TF-IDF embedding using the *TfidfTransformer* class from Scikit-Learn. ii) We used the singular value decomposition (SVD) using the *TruncatedSVD* class to embed the data into 25 components. iii) We transformed the 25 components with the Harmony algorithm correcting for the library ID. iv) We used the UMAP class from the *umap* Python library with the ‘*correlation*’ as a metric, 2.0 as *repulsion_strength*, and 0.01 as *min_dist*. Our Cicero workflow consisted of the following steps: a) load the sparse matrix and creating a Cicero data object with *make_atac_cds* and *binarize* set as *TRUE*, b) aggregate the raw count data with *make_cicero_cds* and *k=50*, where cell similarity used to bin cells is determined in the harmony-transformed UMAP embedding, c) estimate the distance parameter with *estimate_distance_parameter* using *window=250000*, *maxit=100, sample_num=100,* and *distance_constraint=25000* and computed the mean of the distance parameters, d) generate the Cicero models with *generate_cicero_models* using the mean distance parameter and *window=250000*, and e) assemble the connections with *assemble_connections* and save all the Cicero connections.

#### Signac

The Signac methodology to identify enhancers significantly correlated with the expression of a given gene was described in the Method *Peak-to-gene* section of the published study (Stuart *et al*., 2021). The Signac methodology was described in four steps: i) Compute the Pearson correlation between the gene expression and the accessibility of each peak within 500kb. ii) For each peak, compute a background distribution using 200 random peaks matching the GC content, the accessibility, and the sequence length of the peak, and identified with the *MatchRegionStats* R function (https://github.com/stuart-lab/signac/blob/HEAD/R/utilities.R). iii) Compute a z-test using the z-score obtained with the mean and variance of the background distribution. iv) Retain links with *p*-value < 0.05 and |PearsonScore| > 0.05. In order to process the same simulated matrix as the other methods, we reimplemented these four steps in a Python method: a) We first used a 250kb window instead of 500kb to be consistent with our chosen simulation setup and computed the Pearson correlation using the *correlation* function of the Python *NumPy* library. b) In order to randomly select a set of enhancers *E*_*r*_ with similar accessibility to a given enhancer *e*_*ref*_ and its accessibility *a*_*ref*_. We then computed the probability *P*(*e*|*e*_*ref*_) of each *e* ∈ *E* and their corresponding accessibility mean *a*_*e*_ to be in *E*_*r*_ using the normed kernel function and *sigma=50*:

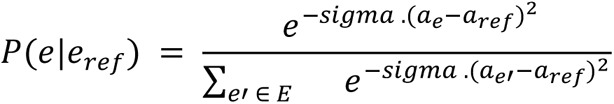

We randomly selected 200 enhancers using these probabilities and computed the Pearson score of each *e* ∈ *E*_*r*_. All the peaks used in the simulation had the same length. c) We transformed the Pearson correlations (of enhancers in the random set *Er* or the query enhancer *e* with the target gene expression) into a *p*-value using the cumulative distribution function *norm*.*cdf* from the *Scipy.stats* package, and excluded links with |Pearson| < 0.05 or *p*-value > 0.05.

#### SnapATAC

The SnapATAC methodology was described in the methods section of the SnapATAC paper (Fang *et al*., 2021). It consisted in considering the expression of a gene *g* as a variable in a univariate logistic regression model that predicted the binary accessibility status of each enhancer within a 1Mb window flanking *g*. For each enhancer *e* from the set of the flanking enhancers *E,* the method built a univariate logistic model *e* = *Logit(g)* using the *glm* function with *link=’binomial’* from the *R* software and used a *p*-value cutoff of 5e-8. We reimplemented the workflow of SnapATAC in a Python function using the *Logit* class with its *fit* method from the *statsmodels* Python library (Seabold and Perktold, 2010). In the original study, a label-transfer procedure was conducted in order to identify the matching cells between two separate snATAC-seq and scRNA-seq datasets. However, this procedure was not necessary in our case because we derived the simulated matrices from a multi-omic snRNA-/snATAC-seq dataset for which we had a matching cell ID between the two modalities.

#### ArchR

The ArchR methodology to infer enhancer-promoter links from scATAC-seq and enhancer-gene correlations from single-cell multi-omic data was described in the supplementary methods of the original study (Granja *et al*., 2021). The method first created a *low-overlapping aggregate of cells* which excludes any pair of cells within this aggregate that share more than 80% of accessible peaks. This was done by first computing a K-nearest neighbor clustering (*K=100* with the Euclidean distance) for a subset of 500 random reference cells and using a 2D embedding of the cells as input. The method iteratively removed cells having >80% of similarities with regards to the KNN neighbors and aggregated the cell vectors of the remaining cells based on their KNN neighbors, with the values further scaled and log-transformed. The Pearson correlation was then computed between the target enhancer and promoter accessibilities, and a *p*-value was inferred by a) transforming the correlation score into a t-statistic: 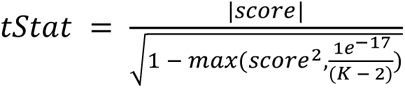 and using the cumulative distribution function of the Student’s *t* law to convert *tStat* into a *p*-value. Finally, an FDR correction was applied using the Benjamini-Hochberg procedure. The R source code of the peak*addCoaccessibility* method is available here: https://rdrr.io/github/GreenleafLab/ArchR/src/R/IntegrativeAnalysis.R and the C++ source code of the iterative aggregation strategy is available here: https://github.com/GreenleafLab/ArchR/blob/master/src/KNN_Utils.cpp. In order to process the same matrices in the same conditions as the other methods, we reimplemented the ArchR strategy in a Python script using the following modifications: a) We used a Harmony + UMAP embedding instead of the iterative LSI strategy used in the original ArchR workflow. The embedding process was the same as described for Cicero. b) We reimplemented the aggregate strategy from the R and the C++ scripts and used the *NearestNeighbors* from the *Scikit-Learn* Python library class with the *cosine* distance to compute the K-neighbors, and iteratively computed the Jaccard similarity to exclude cells from the aggregates. We used the *pairwise_distances* function from Scikit-learn to compute the Pearson scores and the *fdrcorrection* function from the *Statsmodels* library to compute the FDR.

### Intersecting the inferred links with the PCHi-C data

The PCHi-C data are available on GEO (see **Availability of data and materials**) with a limited access using GSE214107 as accession ID. We downloaded the *ibed* files, which are similar to *bedpe* files with the 6 first columns referring to two genomic locations, for each strain, merged them and retained only the unique links.

### Estimating accuracy with the PCHi-C data

We used the physical enhancer-promoter interactions from the PCHi-C datasets from the islet and adipose tissues as references to compute the overall accuracy scores (precision, recall, f1-score) of the links inferred with the Enlink, Cicero, and Chi2+FDR workflows. For both the islet and adipose, we analyzed each cell population independently with each workflow. We then combined all the links obtained for a given dataset and workflow and estimated the overall precision, recall, and f1-score using the set of the PCHi-C links as true positives. We used the *enhtools* software with the *intersect3* option from the Enhlink software suite (see above) to find the intersecting links between the reference set and the results of the different workflows.

### Estimating batch effect with entropy measurements

We quantified the impact of technical batch effects by first reasoning that a true link should be distributed over sequencing batches (that are otherwise biologically similar). We used the information theory principle and computed the Shannon entropy of the link with regard to the sequencing library ID of the dataset. The library ID of the islet and adipose datasets were the ID of the 10X Genomics sequencing runs and were regarded as variables associated with the batch effect. We first computed for each inferred link from an enhancer *e* to a gene *g* the link vector *v*_*l*_ such as

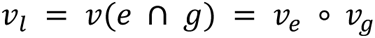

We then computed the Shanon entropy of *v*_*l*_ with regard to the set of library IDs *B*:

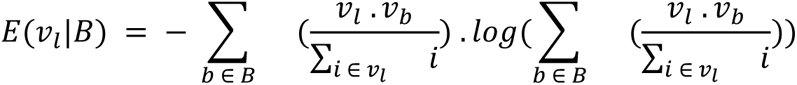

Here, *v*_*b*_corresponds to a binary vector indicating which cells belong to *b*. Finally, we plotted the batch effect entropy distribution using the *seaborn* Python library.

### Inferring Enhlinks atlases for the islet and adipose tissues

For each tissue, we processed each cell type independently using the library ID, sex, diet, and genotype/strain as covariates. We used 0.01 as the *p*-value cutoff, with the *secondOrder* and the *uniformSampling* options to infer the covariate-specific linkages from a uniform distribution of the covariates within each bootstrap sample. For each bootstrap sample, we used a random subset of 66% of the features, a maximum of 4 explanatory features, a depth of 2, and a downsampling size of 15,000 cells. We created a binary cell x promoter sparse matrix by considering all the promoter regions of each gene and give the value 1 for a given cell if at least one read is found, zero otherwise. This matrix is then used as *M*_*target*_ (see **Additional file S1**). We used the *ATACMatUtils* command with the *-use_symbol* option from the *ATACdemultiplex* package (https://gitlab.com/Grouumf/ATACdemultiplex) to create the matrix from the BED file containing the reads and barcode IDs. We then intersected the obtained linkage of each cell type with the PCHi-C links of either the adipose or the islet tissues using the *enhtools* software with the *-intersect3* option

### Inferring Enhlinks for the striatum tissue

We restricted the Enhlink analysis to the two neuron cell types expressing either *Drd1* or *Drd2*. We first performed two Enhlink analyses with the default parameters on the two neuron subtypes expressing either *Drd1* or *Drd2* using a) the scATAC-seq matrix alone and b) the scATAC-seq matrix for the enhancer features and the scRNA-seq matrix to infer enhancers linked to gene expression. We used the default parameters with the library ID as a covariate when using the scATAC-seq matrix only but extended the *max_features* and *depth* parameters to 6 and 4, respectively. We then intersected the ATAC and ATAC+RNA linkages of the *Drd1* and *Drd2* cell type using the *enhtools* executable with the *-intersect* option. In a second analysis aimed at identifying Drd1- or Drd2-specific linkages, we processed with Enhlink all the cells from either the *Drd1* or the *Drd2* neurons and used the library ID together with the cell type (Drd1 or Drd2) as covariates. We performed two processing steps using either the ATAC-seq data alone or the ATAC-seq data combined with the RNA-seq data that we further intersected with *enhtools*. We also processed the same ATAC-seqs dataset with Cicero using the protocol described above.

### Preparing eQTL collections

Bulk RNA-sequencing for eQTL analysis was accessed and downloaded from the Churchill Lab QTL viewer (https://qtlviewer.jax.org/viewer/CheslerStriatum accessed 02/17/2023). This dataset represents striatum samples collected from individual mice (n = 368) from the Diversity Outbred genetic reference population (The Jackson Laboratory catalog #009376). The downloaded package includes a gene expression estimate matrix, genotype probabilities, kinship matrix and metadata including sex. We performed eQTL mapping restricted to the set of genes differentially expressed between *Drd1* and *Drd2* neurons (n = 96 and 103 respectively). eQTL mapping was performed using a linear mixed model to account for kinship on normalized, transformed gene-level expression values using the ‘scan1’ function in r/qtl2 (Broman *et al*., 2019), including sex as an additive covariate. Genes passing a filter requiring a local-eQTL with a LOD score greater than 8 (n = 162) were further included for SNP association using the ‘scan1snps’ function in r/qtl2. All variants within a 1.5 LOD confidence interval were included for association analysis. Resulting variants with LOD score drop of 1.5 from the maximum were retained, along with their strain distribution pattern, to intersect with Enlink significant links and compare to haplotype effect pattern for the linked eQTL gene. The haplotype effects of the QTL for the candidate genes *Kcnb2*, *Gulp1*, and *Col25a1* were estimated using ‘scan1blup’ from r/qtl2.

### Intersecting Enhlinks from the striatum with eQTL

We formatted the results of the eQTL analysis described above as a BED file listing the genomic coordinates of the SNPs and intersected these regions with the enhancers for the Drd1 and Drd2 neuron subtypes that were correlated to both promoter accessibility and expression of their corresponding genes (see above). We then computed a *p*-value using the Mann-Withney procedure to test if the overlapping enhancer presented differential accessibility between the genotypes. We used the *p*-values and the LOD score to identify the enhancers of *Kcnb2*, *Gulp1*, and *Col25a1*. For each enhancer and their associated promoter and gene, we computed the barplot of accessibility (for the enhancer and promoter) or gene expression, using the four 10X Genomics libraries as individual measurements.

### Evaluating Enhlink and Cicero with the striatum dataset

We intersected the linkages obtained from Cicero and Enlink for the Drd1/Drd2 neuron cell types from the ATAC-seq data with the 96 and 103 marker genes of the Drd1 and Drd2 neurons, respectively. For each gene and its inferred enhancer, we computed a univariate logistic regression: *Ve = f(Vg)*, with Ve the boolean vector of the enhancer *e* and of size *1 x cell*, indicating if *e* is accessible for each cell, and *Vg* the numerical vector indicating the scaled gene expression of *g* for each cell. We used the *Logit* class with its *fit* function from the Python *statsmodels* library to infer the *p*-value of each model. We then compared the *-log10(p-value)* distribution of the enhancer from Enhlink and from Cicero for different cutoffs: 0.0, 0.1, 0.2, and 0.28. We chose 0.28 as a cutoff to obtain a number of links (704) similar to the number obtained with Enhlink (802). We used the Mann-Whitney test to assess the difference of the *p*-value distributions between the links from Enhlink and with Cicero results. Since no difference was observed between Enhlink and Cicero with 0.28 as cutoff, we applied a t-test in this case, assuming normality of the distributions.

## Supplementary materials

**Figure S1** f1-score, precision, and recall computed from scATAC-seq data and compared to their corresponding random score distribution using the cell types aCM, vCM, or all the cells for one enhancer from *KCNH2* and three enhancers from *MYL2*.

**Figure S2** Detailed Enhlink analytical workflow. Inputs are: *M*_*peak*_ the cell x peak matrix, *M*_*target*_ and *M*_*cov*_ (optional), the cell x target and cell x covariate matrices, *G* the list of target regions, and *Cl* (optional) a cell clustering. If *Cl* is not provided all the cells will be assigned to one cluster. *M*_*i*_ and *M*′_*i*_ are the matrices from which the significant features associated with the target vectors are extracted. The random subsampling of the samples and features done on these matrices are not represented here for simplicity. *v*_*feature*_, *v*_*target*_, and *v*_*eg*_ are cell vectors indicating the accessibility of a region or the expression of a target. *Cov*′ is a set of significant covariates associated with *v*_*target*_. The dashed procedures are optional.

**Figure S3** Importance of hyperparameters on Enhlink accuracy. *dataset* refers to the type of dataset used, *downsample* refers to the number of cells randomly selected for each tree and is the most important hyperparameter with regards to accuracy (higher is more accurate), *max_features* refers to the maximum number of explanatory features used and has low importance with regards to accuracy, the other variable names in x axis refer to Enhlink hyperparameters described in **Table S1**.

**Figure S4** f1-score, precision and recall on 897 simulated genes and 4090 simulated enhancers when using different low-overlapping aggregate parameters of the ArchR method. *K* refers to the number of neighbors of the KNN steps (100 by default) and NS refers to the number of random samples used as references (500 by default). The result highlighted in purple refers to the default hyperparameter values used by ArchR. The Signac approach was used to infer a *p*-value from a random distribution.

**Figure S5** Histogram distribution of the *batch x link* entropy for Cicero, Chi2 and Enhlink and Maximum entropy value according to the number of classes used. The entropy is maximum when all the classes are equiprobable. 18 classes corresponding to the 18 libraries were used to compute the covariates entropy.

**Figure S6** Number of links per tissue (islet or adipose) and cell type, annotated according to overlap with PCHi-C and by biological covariate.

**Figure S7 A** UMAP projection of the single-cells and clusters for the snRNA-seq Striatum dataset. Enhlink score (**B**) and *p*-value (**C)** distributions of links inferred from promoter accessibility in *Drd1* and *Drd2* neurons separated by whether they were concordant or not with links inferred from gene expression.

**Figure S8** Comparison of enhancer-gene expression association between Cicero and Enhlink links inferred from the multi-omic striatum dataset when using only the snATAC-seq data to infer enhancers. **A** Number of links inferred for the marker genes of *Drd1*/*Drd2* neurons by each method and for different Cicero cutoffs. **B** Number of genes inferred for each method and cutoff. **C** Distribution of the -log(*p*-value) of the univariate logistic regression using enhancer accessibility as dependent variable and gene expression as response.

**Figure S9** Chromatin accessibility of distal *Drd1*-associated enhancers. **A** Four OCR regions linked to the top 10 striatonigral marker genes, including *Drd1* (grey). In addition, three *cis*-OCR regions are also linked to the *Drd1* promoter (green). **B** Exonic region (in grey) of *Isl1* correlated with both promoter accessibility and expression of nine out of the top 10 striatonigral (expressing *Drd1*) marker genes. A *cis*-OCR region linked to *Isl1* promoter is highlighted in green.

**Table S1** Description of Enhlink’s hyperparameters

**Table S2** Description of the hyperparameter values used for the simulation experiment

**Additional file S1** Extended description of the Enhlink procedure

## Funding

Research reported in this publication was supported by The Jackson Laboratory Cube Initiative and the Jackson Laboratory *Scientific Support Internal Fund* (SSIF) mechanism with the project id: 19005-21-05. Further funding was provided, in part, by the National Institute of General Medical Sciences grant R35GM133724 to CLB and the Jackson Laboratory Director’s Innovation Fund to CLB and EJC. P50 DA039841 supports the Center for Systems Neurogenetics of Addiction to EJC. Data generation through the Single Cell Biology service was supported in part by the JAX Cancer Center (P30 CA034196). Additional support was also provided by The Jackson Laboratory Cube Initiative.

## Author contributions

OP and BSW envisioned this project. OP developed Enhlink and implemented the project. OP, WZ, DS, and CLB conducted the analyses. CLB performed the eQTL analysis. OP, BSW, and CLB wrote the manuscript with inputs from all authors. EJC supervised the eQTL collection analysis. CLB supervised the striatum tissue collection and experimental design. CS and AO performed the tissue collection. SD performed the single-cell multiome experiment.

## Acknowledgments

We gratefully acknowledge the contribution of the Single Cell Biology service, the Genome Technologies service, and Cyberinfrastructure high performance computing resources at The Jackson Laboratory for expert assistance the work described herein. These shared services are supported in part by the JAX Cancer Center (P30 CA034196). We would like to also thank Dr. Mike Lloyd and Dr. Anuj Srivastava for their help in obtaining and processing the PCHi-C data, and Dr. Vivek M. Philip, Dr. Robyn Ball and Leona Gagnon for managing and executing experimental designs.

## Availability of data and materials

The PCHi-C data used in this manuscript are currently available with limited access on request. The PCHI-C data are planned to be released to the public on December 31, 2023. In the meantime, these data will be available on request. The multi-omics striatum dataset is publicly accessible using the following accession number: <u>GSE228530</u>. The Linkage atlases for the islet and adipose datasets are available as Figshare datasets with the following DOIs: https://doi.org/10.6084/m9.figshare.22336033.v1 (adipose) and https://doi.org/10.6084/m9.figshare.22335919.v1 (islet).

## Competing interests

The author(s) declare(s) that they have no competing interests.

## Notes

https://gitlab.com/Grouumf/enhlinktools

https://enhlinktools.readthedocs.io/en/latest/

### Competing Interest Statement

The authors have declared no competing interest.

