## Supplementary file S1: Extended methods for "Enhlink infers distal and context-specific enhancer-promoter linkages"

### Enhlink detailed procedures

The analytical procedure performed by Enhlink consists of several steps schematized in **Figure S1.** It can be summarised as follow:

a. For each target genomic region, create a feature matrix and a response vector
b. Model the response vector as a function of the feature matrix and find the significant features linked to the target region.
c. Optionally perform a second order analysis to identify the specific covariates associated to the links.

#### Input matrices

Enhlink requires a sparse matrix $M_{peak}$ of shape $cell\,x\,features$ for the set of features (i.e., OCRs or peaks) *P*. The second mandatory input is a list of genomic target regions *G*, defined in a *Browser Extensible Data* (BED) file with an ID (e.g. gene symbol) annotating each region. As an example, the genomic region referring to one of the mouse mm10 *Kcnh5* promoter regions would be defined by: *“chr5 24341900 24345900 Kcnh2”*.

Optional inputs are

a) a *cell x target* matrix $M_{target}$. The targets are promoter or locus accessibilities (e.g., when analyzing scATAC-seq data only) or gene expression (e.g., when analyzing multi-omic snATAC/RNA-seq data). $M_{target}$ for locus accessibilities can for example be useful to agglomerate the read count of the gene bodies or when the read count of multiple promoter locations of a each gene are used.
b) a *cell x cov* covariate matrix $M_{cov}$.
c) cell clusters *Cl* defined in a two-column (cellID, clusterID) TSV file.

$M_{peak}$ and $M_{target}$ should be provided as sparse matrix files, using the Matrix Market (MTX) format, the Cell-Ranger MTX format, or the Coordinate List (COO) format. The clusters and the covariates should be provided as TSV files. The clusters should be in a two-column file with the first column being the cell IDs and the second the cluster IDs.

#### Response vectors

Enhlink iterates through the regions of G, and for each genomic region $g \in G$, it constructs a *cell x n* feature matrix $M_n$ with *n* the number of *p* features from *P* such as $p \in g'$ and $p \notin g$. *g'* is defined as an extended interval of *g*: $g' = \{g_{min} - r, g_{max} + r\}$, with *r* a hyperparameter set by default at 250kb. Note that if *r* is set to 0, then the full set of *P* will be used as features of $M_n$. The minimum number of neighborhood features is defined by a hyperparameter h set at 100 by default. If n < h, Enhlink creates $h - n$ random new features by a) randomly sampling with replacement *h - n* features within the *n* features and b) shuffling each randomly sampled feature. If $M_{cov}$ is provided, each covariate is one-hot encoded with the first category of each feature removed. Enhlink then creates a response vector for *g,* merging either the feature vectors $v_{peak}$ from $M_{peak}$ such as $p \in g$ or, if $M_{target}$ is provided, it extracts $v_{target}$ from $M_{target}$ based on the matching ID of *g* with the feature index of $M_{target}$.

If *Cl* is defined, Enhlink iterates through *G* and *Cl,* and for each cluster $C \in Cl$, Enhlink uses the cell contained in *C* as a cell index rather than all the cells. The instantiation of $M_n$, $v_{target}$ can be summarized by the following procedure:

Function Enhlink(minSize h):

1. Load $M_{peak}$, *G* from files
2. If provided, load $M_{target}$, $M_{cov}$, *Cl* from files
3. If *Cl* not provided, all cells belong to the same cluster.

4. For each region g from G do
   a. $M_n$, *n* = constructNeighborhoodMat($M_{target}$, *g*)
   b. If *n* > *h* { $M_n$ = addRandomFeatures($M_n$, h -n) }
   c. If $M_{cov}$ exists { $M_n$ = {$M_n$, $M_{cov}$} }
   d. $v_{target}$ = defineVtarget($M_{peak}$, $M_{target}$, *g*)
   e. For *C* in *Cl* do
      i. ScoreFeaturesForCluster($M_n$, $v_{target}$, *C*)

**Procedure 1** Enhlink overview with the input matrix $M_{peak}$, the set of genomic region G, the target matrix $M_{target}$, the covariate matrix $M_{cov}$, the cell clusters *Cl*, the feature matrix $M_n$, the number of neighbor features n, the minimal number of features h, and the response vector $v_{target}$. ScoreFeaturesForCluster subsets $M_{peak}$ and $v_{target}$ according to the cells of cluster *C* and identify significant features within the cells of cluster *C*.

#### Feature scores computation

For each genomic region $g$, Enhlink finds the features significantly predictive of $v_{target}$, the binary response vector of $g$. $v_{target}$ is modelled as a function of the features from the feature matrix $M_n$ and the covariates:

$$v_{target} = f_{Enhlink}(M_n, M_{cov})$$

##### Bootstrapping

More precisely, $M_n$ and $M_{cov}$, the covariate matrix, are stacked together into $M_i$ a matrix with Y columns. Enhlink follows a strategy similar to a random forest classifier (Breiman 2001) to find the best features amongst $M_i$ that predicts $v_{target}$. The workflow performs *k* iterations with *k* a hyperparameter set by default at 100. For each iteration, Enhlink selects a random subset of features F from Y according to the hyperparameter *m* (by default it uses all the features) and creates a bootstrap N' (i.e. resampling with replacement), set by default as |N'|=|$M_i$|. The process can be summarized by the following procedure:

Function ScoreFeatures(sample N, feature Y, depth d, nbFeat m, nbCells n):

1. Define scoreDictAll as map(featureKey -> floatArray)
2. For each k iteration do
   a. F = drawRandomFeatures(Y, m)
   b. N' = drawBootstrapSample(N, n)
   c. scoreDict = ReccursiveFeatureScoring(N', F, d)
   d. Update scoreDictAll with scoreDict
3. ComputePValues(scoreDictAll)

**Procedure 2** Enhlink feature scoring overview taking as input a set of cells *N*, a input set of features *Y*, a number of cells to draw with replacement *n*, a number of features to draw *m*, and a depth in the decision tree d. *N'* and *F* are a random subset of cells with from *N* and *Y*. scoreDict is a mapping of a feature to a score.

##### Decision trees

For each iteration, Enhlink follows a strategy similar to a Decision tree (Quinlan 1986) by computing a modified Information Gain score for each feature from *F* and by recursively reducing the number of samples and features according to the top feature selected at each level. The recursive procedure can be summarized with the following procedure:

Function ReccursiveFeatureScoring(samples N', features F, depth d, scoreDict D):

1. Find $f_{max} \in F$ for which $score = IG'(f_{max})$ is maximal
2. Remove $F_{max}$ from F: $F' = F - f_{max}$
3. Split N' into two subsets: N0, N1 based of $f_{max}$ value for each $n \in N$
4. d++
5. $D[f_{max}] = score$
6. check if conditions are met for return
7. if|N0| > minLeafSize { ReccursiveFeatureScoring(N0, F', d, D)}
8. if|N1| > minLeafSize { ReccursiveFeatureScoring(N1, F', d, D)}

**Procedure 3** Enhlink recursive feature scoring Enhlink feature scoring overview taking as input a set of cells N', a input set of features F, depth in the decision tree d, a feature to score mapping scoreDict. minLeadSize is a fixed value (10 by default) indicating the minimum number of sample required to keep performing the recursive procedure. $f_{max}$ indicates the feature from F with the maximal score.

The conditions for which *ReccursiveFeatureScoring* is returning are a) $f_{max}$ = 0, b) *d* >= $d_{max}$ with $d_{max}$ defined as a hyperparameter and set to 2 in the current version c) *F'* is empty d) the number of features processed |*D*| is larger or equal to *nbFmax*, a hyperparameter set to 4 in the current version and e) if both N0 and N1 have fewer elements than minLeafSize, a hyperparameter set to 10 by default.

#### Modified Information Gain score

Given the corresponding feature vector $v_{feature}$, the Information Gain defined by the formula below is first computed:

$$IG(f) = E(v_{target}) - ((\frac{|v_{feature} == 1|}{|v_{feature}|}) . E(v_{target} | v_{feature} == 1) + \frac{|v_{feature} == 0|}{|v_{feature}|} . E(v_{target} | v_{feature} == 0))$$

with E, the entropy defined for a binary vector by:

$$E(v) = \frac{|v == 0|}{|v|} . log(\frac{|v == 0|}{|v|}) + \frac{|v == 1|}{|v|} . log(\frac{|v == 1|}{|v|})$$

The notation $v_{feature} == 1$ refers to the vector extracted from $v_{feature}$ for which all elements are equal to 1. The notation $v_{target} | v_{feature} == 1$ refers to the vector extracted from $v_{target}$ for which its elements have an index i such as $v_{feature}[i] == 1$. For each feature, the modified information-gain score *IG'(f)* is computed by scaling *IG(f)* with the ratio of non-null cells from $v_{feature}$ and the ratio of non-null cells from $v_{target}$ to favorite features with high I*G(f)*, and high ratio of accessible enhancers:

$$IG'(v) = IG(v) . \frac{|v_{feature} == 1|}{|v_{feature}|} . \frac{|v_{target} == 1|}{|v_{target}|} . Sign(Pearson(v_{target}, v_{feature}))$$

The $\frac{|v_{target} == 1|}{|v_{target}|}$ part of the equation is identical along all the features of *F* for a given iteration, but changes as the recursive process shrink *N*. $Sign(Pearson(v_{target}, v_{feature}))$ refers to the sign of the Pearson correlation between $v_{target}$ and $v_{feature}$ and indicates whether $v_{target}$ and $v_{feature}$ are correlated or anticorrelated.

#### P-values computation

For a given genomic region *g*, Enhlink performs *k* iterations and for each iteration, it computes and stores the score of the best features *Fs* based on the modified Information Gain *IG'* formula described above. Enhlink then computes the p-value to have an *IG'* score different than 0 for each feature from *Fs.* These features have at least one best *IG'* score from the *k* iterations. For each feature of *Fs*, Enhlink computes the mean and the standard deviation (std) over the k iterations and then computes the t-test p-value using the CDF function of the Go *distuv* package (https://pkg.go.dev/gonum.org/v1/gonum/stat/distuv). It then corrects the p-values using the False Discovery Rate (FDR) procedure of Benjamini-Hochberg.

#### Simulated enhancers / promoter

During the inference of the enhancers of a given genomic region *g*, Enhlink can optionally generate simulated a promoter and linked enhancers specific to *g* and estimate the expected True Positive Rate (TPR), True Negative Rate (TNR), False Positive Rate (FPR), False Negative Rate (FNR) for the given genomic region *g*. The generation of the simulated enhancer-promoter links is controlled by three variables a) the number of simulated links to generate b) the type 1 noise which is controlled by the λ parameter of a first Poisson distribution ($\lambda_{close}$) and c) the type 2 noise which is controlled by the λ parameter of a second Poisson distribution ($\lambda_{open}$). The simulated promoter vector $v^{s}_{target}$ is obtained by simply shuffling the promoter vector $v_{target}$ described above. Each simulated enhancer $v^{s}_{e}$ associated to $v_{target}$ is first instantiated by duplicating $v^{s}_{target}$. Secondly, for each value *v* of $v^{s}_{e}$ a random number *r* is drawn following the Poisson distribution with either $\lambda_{close}$ if v==1 or $\lambda_{open}$ otherwise. If $\lambda_{close}$ is used and *r > 0,* then v is set to 0. If $\lambda_{open}$ is used and *r > 0,* then *v* is set to 1. The simulation process can be summarized by the following procedure:

Function SimulateEnhancersPromoter(matrix $M_n$, region g, nbFeatures, $\lambda_{close}$, $\lambda_{open}$)

1. $v_{target}$ = obtainVectorFromRegion(g)
2. $v^s_{target}$ = shuffle($v_{target}$)
3. Me = EmptyMat()

4. For i in range(1, nbFeatures) do
    a. $v^s_e$ = copy($v^s_{target}$)
    b. $v^s_e$ = addλCloseNoise($v^s_e$, $v^s_{target}$, $\lambda_{close}$)
    c. $v^s_e$ = addλOpenNoise($v^s_e$, $v^s_{target}$, $\lambda_{open}$)
    d. $M^s_e$ = stackVectorToMatrix($M^s_e$, $v_e$)

5. $M^s_n$ = stackMatrixToMatrix($M_n$, $M^s_e$)
6. Return $M^s_n$, $v^s_{target}$

**Procedure 4** Simulating enhancers and promoters taking as input the feature matrix $M_n$, a genomic region g, the number of features to simulate nbFeatures, and two parameters $\lambda_{close}$ and $\lambda_{open}$ controlling the noise. $v_{target}$ refers to the response vector and Me is the matrix of simulated vectors $v_e$ and is empty at he begining. $M^s_n$ is the feature matrix $M_n$ concatenated with Me.

The simulated enhancer matrix $M^s_e$ is stacked with the original feature matrix $M_n$ into a new feature matrix $M^s_n = \{Me, M_n\}$, which contains both real and simulated genomic features. An Enhlink analysis is then performed using $v^s_{target}$ as response vector:

$$v^s_{target} = f_{Enhlink}(M^s_n, M_{cov})$$

The TPR, TNR, FPR, FNR are then inferred from the results of this analysis. The simulation analysis is realized conjointly with the original Enhlink analysis on $v_{target}$ and $M_n$ and the results are stored in separate files. This simulation analysis allows having an estimate of the expected accuracy of Enhlink for a given gene according to the parameters used.

#### Inferring covariate-enhancer-promoter interactions

If the covariate matrix $M_{cov}$ is provided, Enhlink concatenates $M_{cov}$ with the neighborhood features matrix $M_n$ (see **procedure 1**). In this case, Enhlink can optionally infer covariate-enhancer-promoter interactions: for a given genomic region $g$ and its response vector $v_{target}$ the set of covariate features $Cov$ from $M_{cov}$, and the feature matrix $M_n$, Enhlink infers a set of enhancers *E,* significantly co-accessible with *g*, and a set of covariates $Cov'$ correlated with $v_{target}$. For each $e \in E$ *e* and its vector $v_{peak}$ Enhlinks defines $v_{eg} = v_{peak} \circ v_{target}$ as the Hadamard product (e.g. element-wise) between $v_{peak}$ and $v_{target}$. Enhlinks then models $v_{eg}$ as a function of the significant covariates $Cov'$ and $E \setminus e$ (the set *E* without *e*): $v_{eg} = f_{Enhlink}(M_{cov'}, M_{E \setminus e})$

Enhlinks follows the same analytical procedure described above (see *Feature score computation*) to infer the set of covariates $Cov''$ for which a significant interaction between *e*, *g,* and *cov* is detected for all $cov \in Cov'$. The covariate-enhancer-promoter inference can be summarized as follow:

Function getCovEnhancerPromoterInteractions(matrix $M_n$, region *g*)

1. ResultsCov = EmptyLinks()
2. $v_{target}$ = obtainVectorFromRegion(*g*)
3. E, $Cov'$ = getEnhlinkFeatures($M_n$, $v_{target}$)
4. For each e in E do
    a. $M^s_n$ = getMatrixFromFeatureSet(Cv', E \ e)
    b. $v_{eg}$ = HadamardProduct($v_e$, $v_{target}$)
    c. *E'*, $Cov''$ = getEnhlinkFeatures($M^s_n$, $v_{eg}$)
    d. ResultsCov = updateResultsForE(ResultsCov, e, $Cov''$)

**Procedure 5** associating links to covariates taking as input a feature matrix $M_n$ and a genomic region g. $v_{target}$ refers to the response vector, ResultsCov is an empty list of links that is filled incrementally. E refers to the set of enhancers obtained from $M_n$ and $v_{target}$.

#### Addressing Class Imbalance in Samples with Unequal Covariate Distributions

Enhlink optionally offers to address class imbalance in samples with different covariate distributions. This is particularly useful when one or more covariates are over- or under-represented, and can significantly affect the covariate representation in bootstrap samples. Enhlink generates a near-uniform distribution of covariates when the hyperparameter *uniformSampling* is passed as input. The first step in this process is to compute a step length $length$ by dividing the total number of cells by 100. Enhlink then selects the least represented covariate from a count table $t$, chosen randomly at first. Next, $length$ cells are randomly selected from this covariate, and $t$ is updated with the covariates of these cells, since a single cell may have multiple covariates. This process is repeated 100 times to obtain a set of samples with a more balanced distribution of covariates. The procedure can be summarized as follow:

Function generateUnifromSampling(matrix $M_{cov}$, nbSteps int, sampleSize int)

1. covCount = emptyMap()
2. samples = emptyList()
3. *cov* = selectRandomCovariate($M_{cov}$)
4. while |sample| < sampleSize:
   a. For $|M_{cov}|$ / nbSteps:
      i. *cell* = drawRandomCellWithCo($M_{cov}$, *cov*)
      ii. incrementMap(covCount, *cell*)
      iii. updateList(samples, *cell*)
   b. cov = selectLeastRepresentedCo(covCount)

**Procedure 6** generates a uniform distribution of cells based on their covariates. The algorithm takes as inputs the covariate matrix $M_{cov}$, the number of steps nbSteps, and the desired sample size sampleSize. The covariate count covCount is a map that keeps track of how many cells belong to each covariate. Since cells can have multiple covariates, *covCount* is updated accordingly. The algorithm iteratively selects the least represented covariate, randomly selects a fixed number of cells from that covariate, and adds them to the sample list. This process is repeated until the desired sample size is reached.

| **Variable symbol** | **Description** |
|---|---|
| $M_{peak}$ | Matrix of cells x peaks |
| $M_{target}$ | Matrix of cells x target regions |
| $M_{cov}$ | Matrix of cells x covariates |
| $Cov$ | Set of covariates |
| *G* | Set of target regions |
| *P* | Set of peaks from $M_{peak}$ |
| *Cl* | Cell clustering |
| *g* | A genomic region from *G* |
| *r* | Hyperparameter of Enhlink defining a genomic range (250kb by default) |
| *g'* | An extended genomic region of *g* such as $g' = \{g_{min} - r, g_{max} + r\}$ |

| | |
|---|---|
| $M_n$ | Cell x peak matrix with the peaks in the neighborhood of a target genomic region |
| $v_{target}$ | Vector containing the accessibility or expression value of a target region for each cell |
| $v^s_{target}$ | Simulated target vector obtained by shuffling $v_{target}$ |
| $v_{peak}$ | Binary vector indicating if a peak is accessible for each cell |
| $v^s_e$ | Simulated enhancer vector from $M^s_e$ |
| $M^s_e$ | Simulated enhancer matrix |
| $M_i$ | Concatenation of $M_n$ and $M_{cov}$ |
| *Y* | Set of feature columns of $M_i$ |
| *F* | Random set of features from *Y* |
| *E* | Peaks significantly associated with $v_{target}$ |
| *N'* | Random bootstrap samples of the cells of $M_i$ |
| $v_{feature}$ | Binary vector extracted from $M_i$ |
| $v_{eg}$ | Hadamard product between $v_{target}$ and $v_{peak}$ |
| $Cov$' | Covariates significantly associated with $v_{target}$ |
| $cov$ | A covariate from $Cov$ |
| *IG'(x, y)* | Modified Information gain between two vectors x and y |

**Table S1.1** Description of the main variables and symbols used in the extended method.
