## Supplementary material for "Enhlink infers distal and context-specific enhancer-promoter linkages": Table S1

| Name | Description | Type | Default value | Governs speed | Regularization | Drive accuracy* |
|---|---|---|---|---|---|---|
| depth | Max tree depth | Int | 2 | Yes | Yes | No |
| downsample | Max number of samples used for each bootstrap | Int | None | Yes | Yes | Yes |
| keep_sparse | Keep the feature matrix sparse to save memory but significantly slow computation | bool | FALSE | Yes | No | No |
| lambda1 | Controls the amount of dropouts of the simulated variables | Float | 1.2 | No | No | No |
| lambda2 | Controls the amount of False positives of the simulated variables | Float | 0.15 | No | No | No |
| ignoreEnhancerWeight | ignore the ratio of accessibility in the computation of the modified information gain | bool | FALSE | No | No | No |
| neighborhood | define the genomic range for including genomic features surrounding a target region | int | 250000 | Yes | No | No |
| maxFeatType | Maximum number of features to be used for a given tree | Int / Float / str | all | Yes | Yes | Yes |
| max_features | Maximum number of explanatory features for a given tree | Int | 4 | Yes | Yes | No |
| merging_cutoff | Cutoff (bp) for merging close promoter peaks together | Int | 1500 | No | No | No |
| min_leafsize | Minimum size of a tree leaf | Int | 10 | No | Yes | No |
| min_matsize | Minimum matrix size used for computation | Int | 100 | Yes | Yes | No |
| n_boot | Number of bootstrap regression trees to construct for each model | Int | 100 | Yes | Yes | Yes |
| nb_sim_features | if > 0, number of simulated features to generate for each target region to estimate expected accuracy | Int | 0 | No | No | No |
| onlySim | Only perform the simulation analysis by shuffling target region and introducing simulated enhancers. | bool | FALSE | Yes | No | No |

| | | | | | | |
|---|---|---|---|---|---|---|
| secondOrder | Identify the covariates associated with each inferred enhancer-promoter link | bool | FALSE | Yes | No | No |
| secondOrderMaxFeat | Maximum number of features to be used for a given tree for the secondOrder inferance | Int / Float / str | 2 | Yes | Yes | No |
| rmPeaksInPromoter | Exclude from model features which are within promoter boundaries | bool | FALSE | No | No | No |
| threads | Number of internal threads | Int | 2 | Yes | No | No |
| threshold | FDR p-value threshold | Float | 0.05 | No | Yes | No |
| uniformSampling | If true, sample cells to have a uniform distribution of the covariates at each bootstrap | bool | FALSE | No | No | No |

* higher values leads to better results
