## Supplementary material for "Enhlink infers distal and context-specific enhancer-promoter linkages": Table S2

| Parameter name | Values | Description |
|---|---|---|
| maxFeatType | 0.33, 0.66, 1.0 | Ratio of the number of features used |
| downsample | 100, 1000, 5000, 10000 | Number of samples used for each tree |
| n_boot | 40, 100, 150 | number of trees |
| λclose | 1.2, 1.4 | Type 1 noise |
| λopen | 0.15, 0.25 | Type 2 noise |
| max_features | 2,4,8, | Maximum number of explanatory features per tree |
| depth | 3, | Depth of the tree |
| min_matsize | 50, 100, 150 | Min number of features for Mn |
| bb_sim_features | 2, 4, 8, | Number of simulated enhancers to generate |
| Threshold | 0.10, 0.05, 0.01 | p-value cutoff |
