## Supplementary figures and images for "Enhlink infers distal and context-specific enhancer-promoter linkages"

### Figure S1

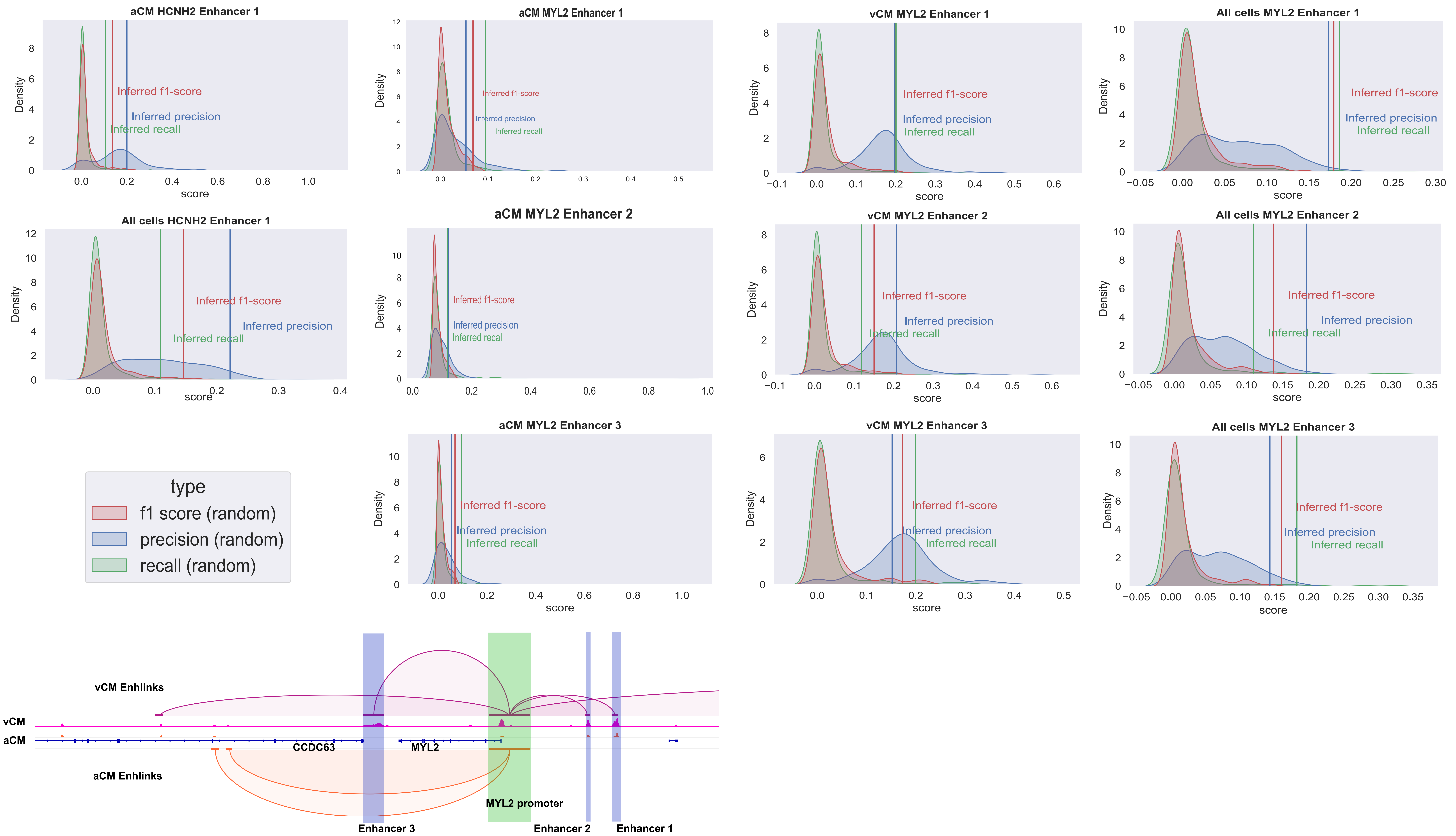

### Figure S2

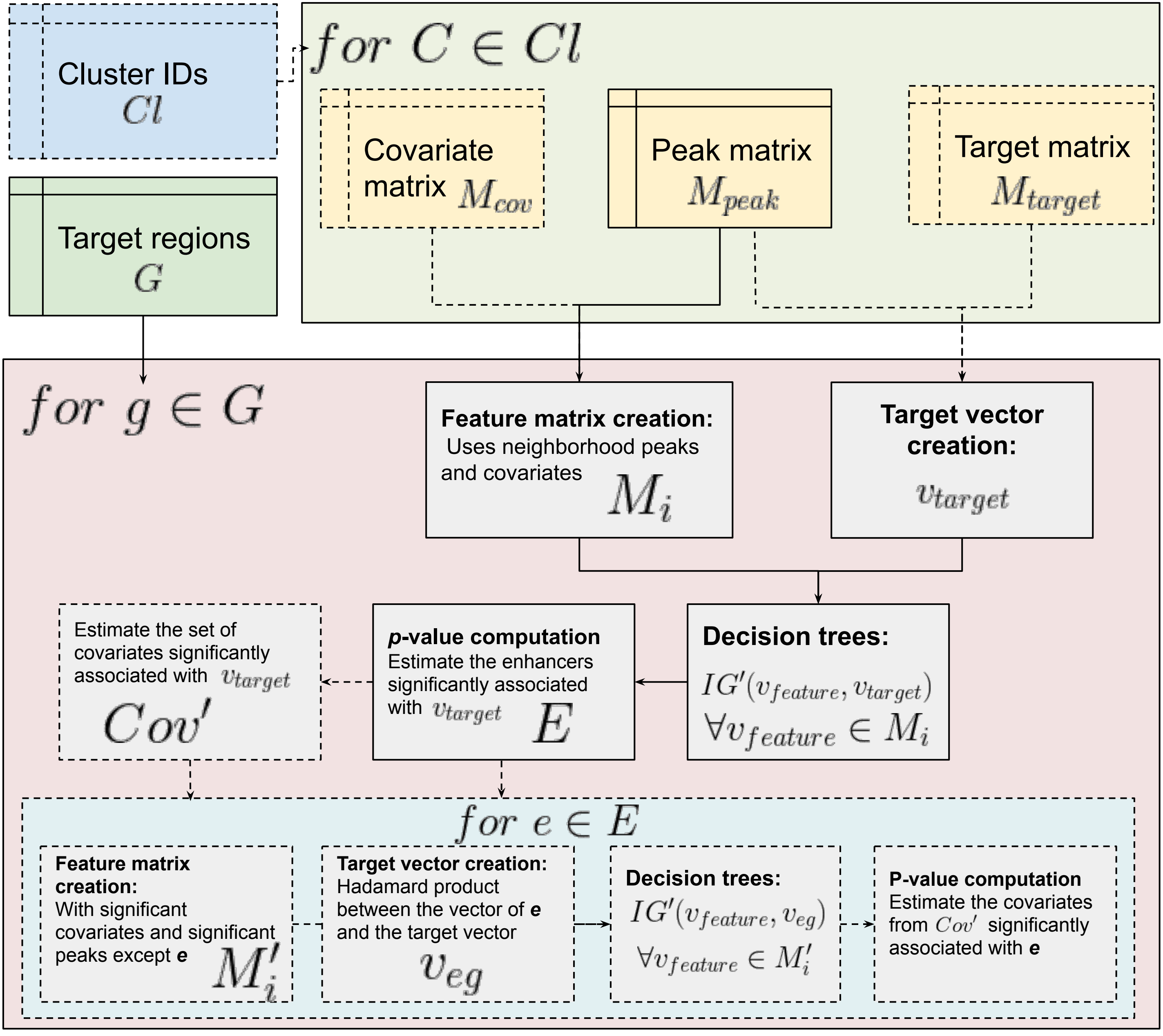

### Figure S3

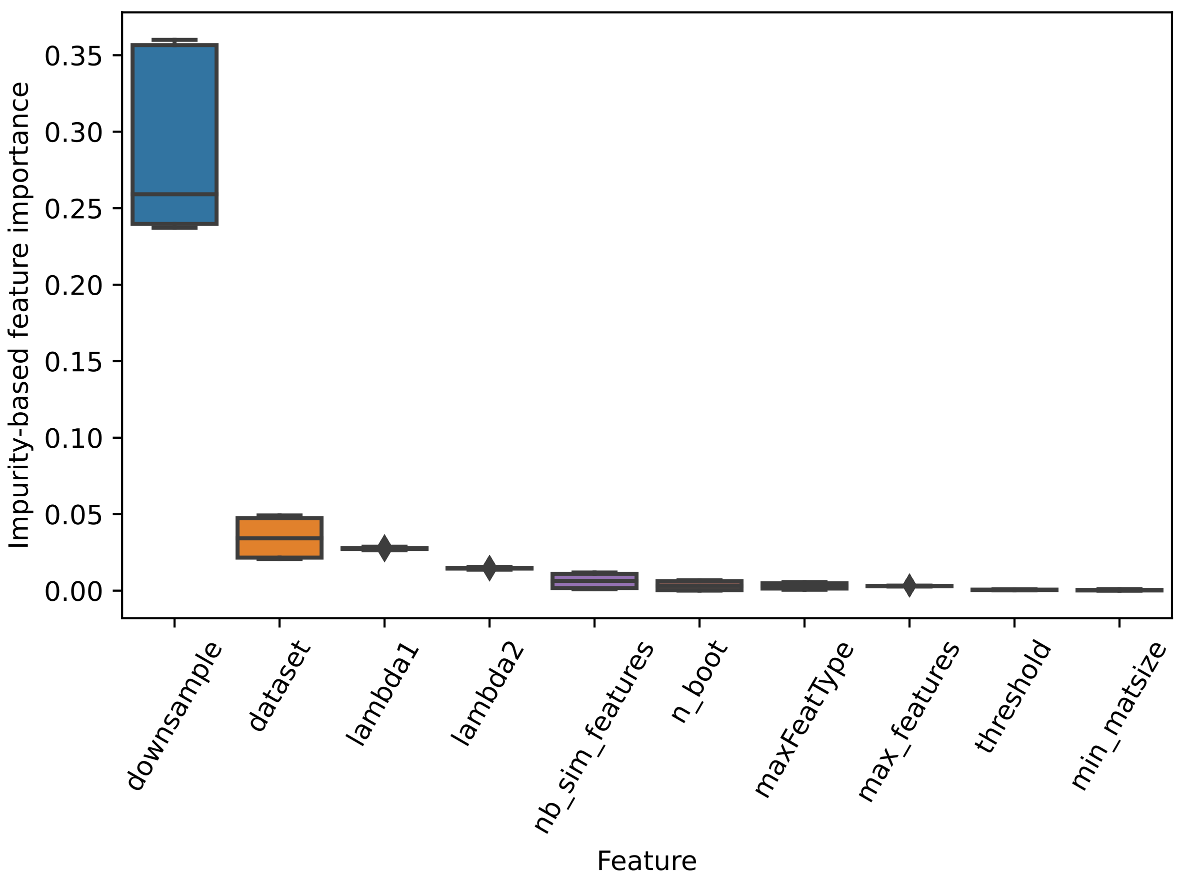

### Figure S4

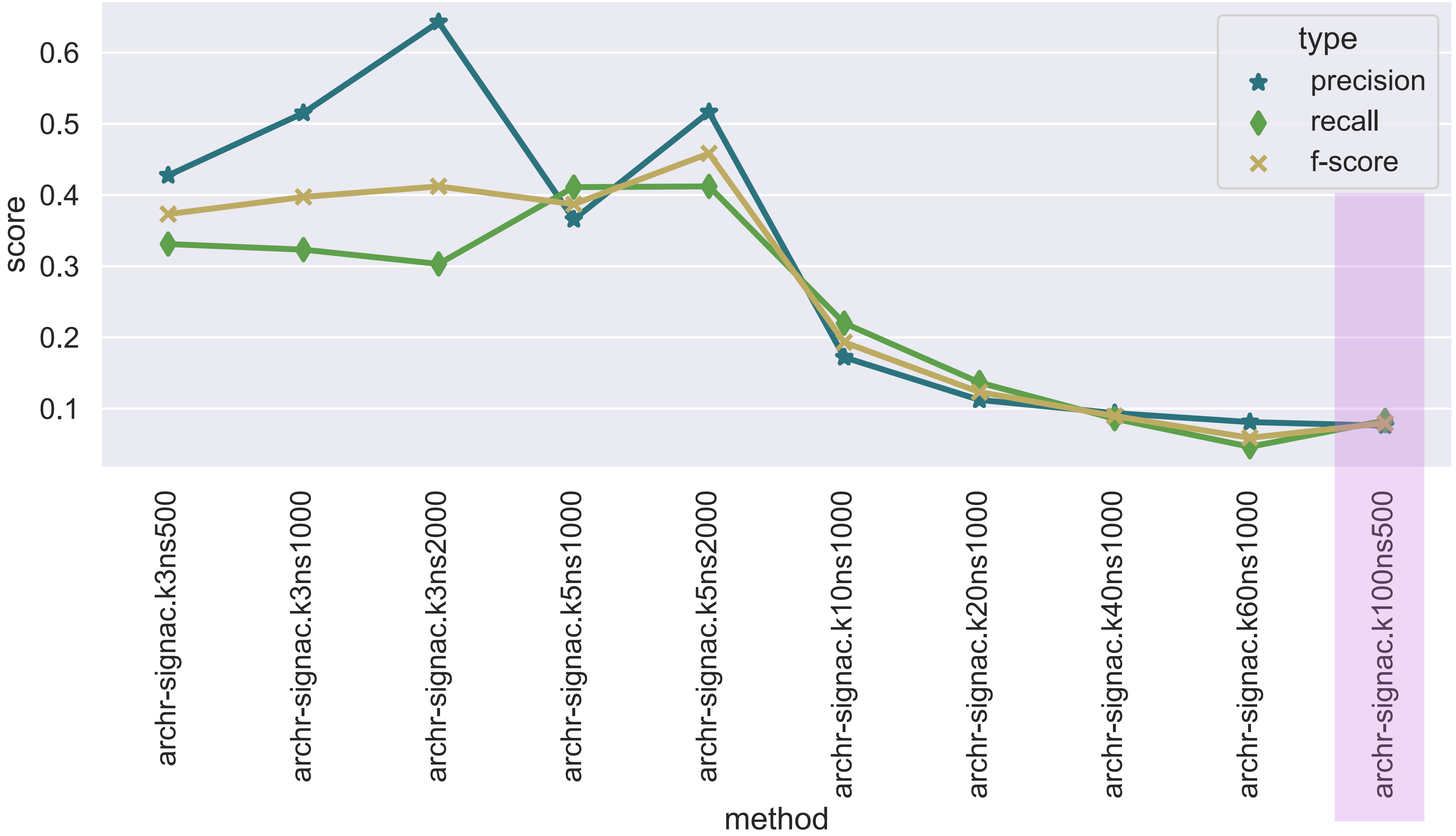

### Figure S5

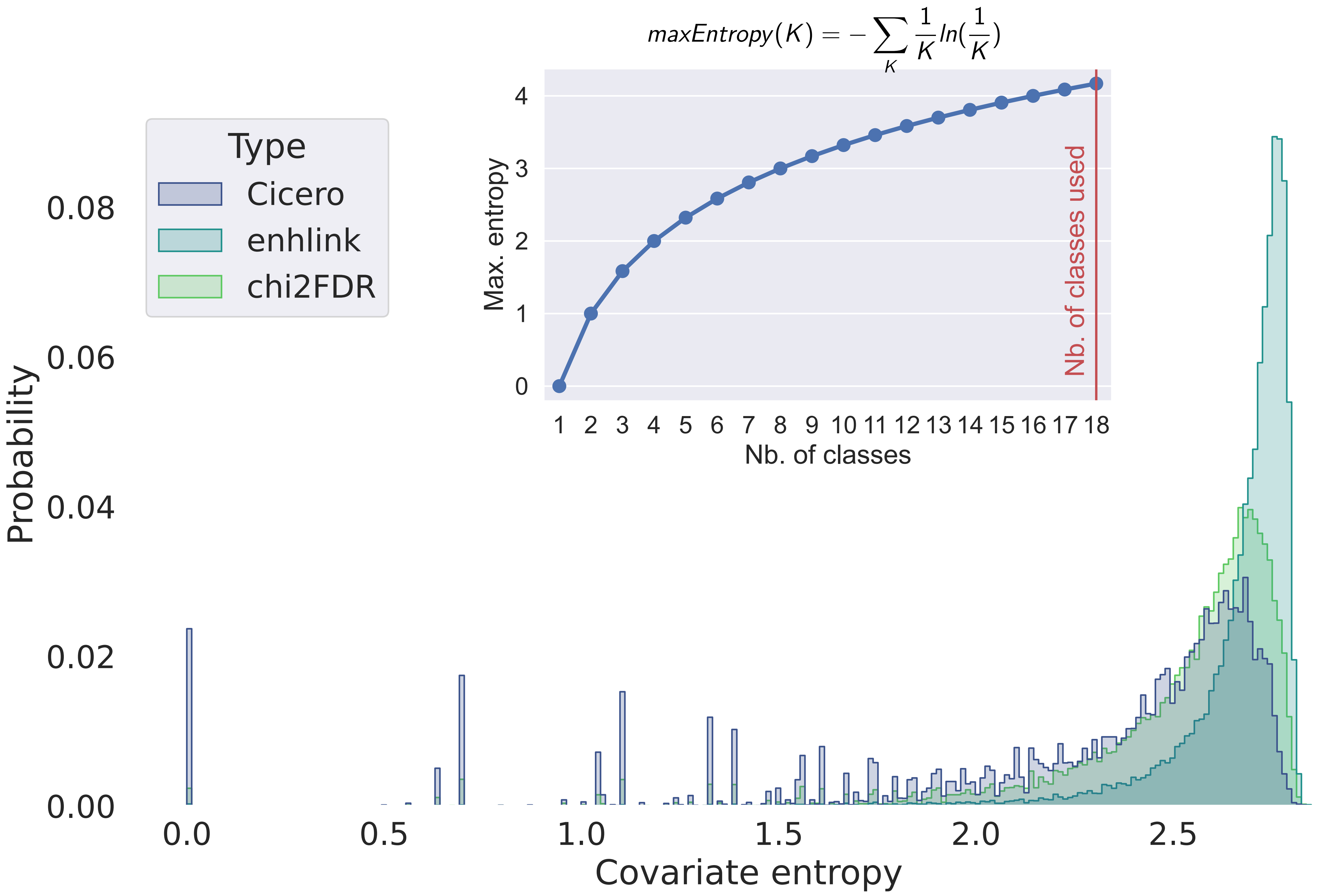

### Figure S6

#### Adipose

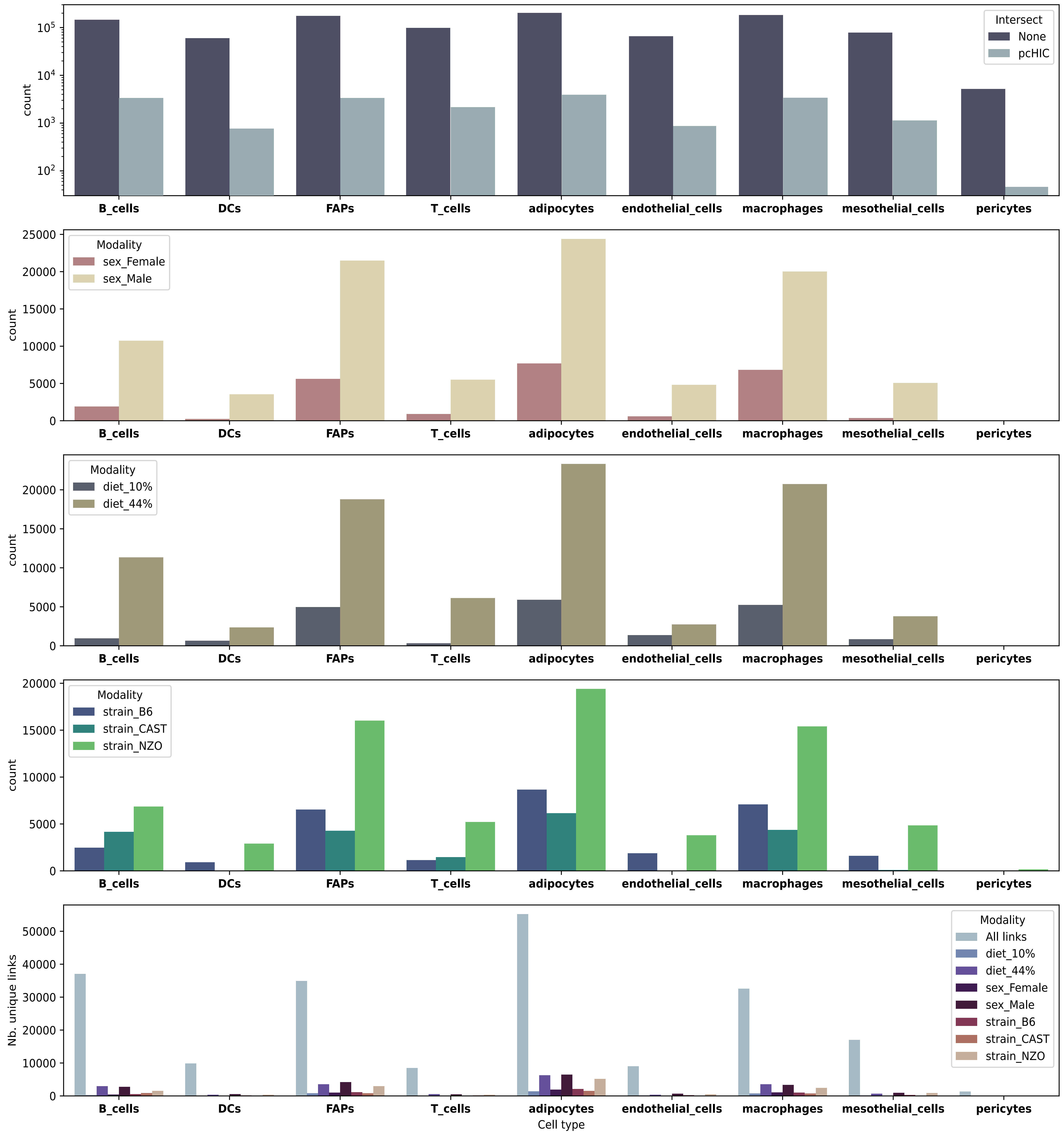


#### Islet

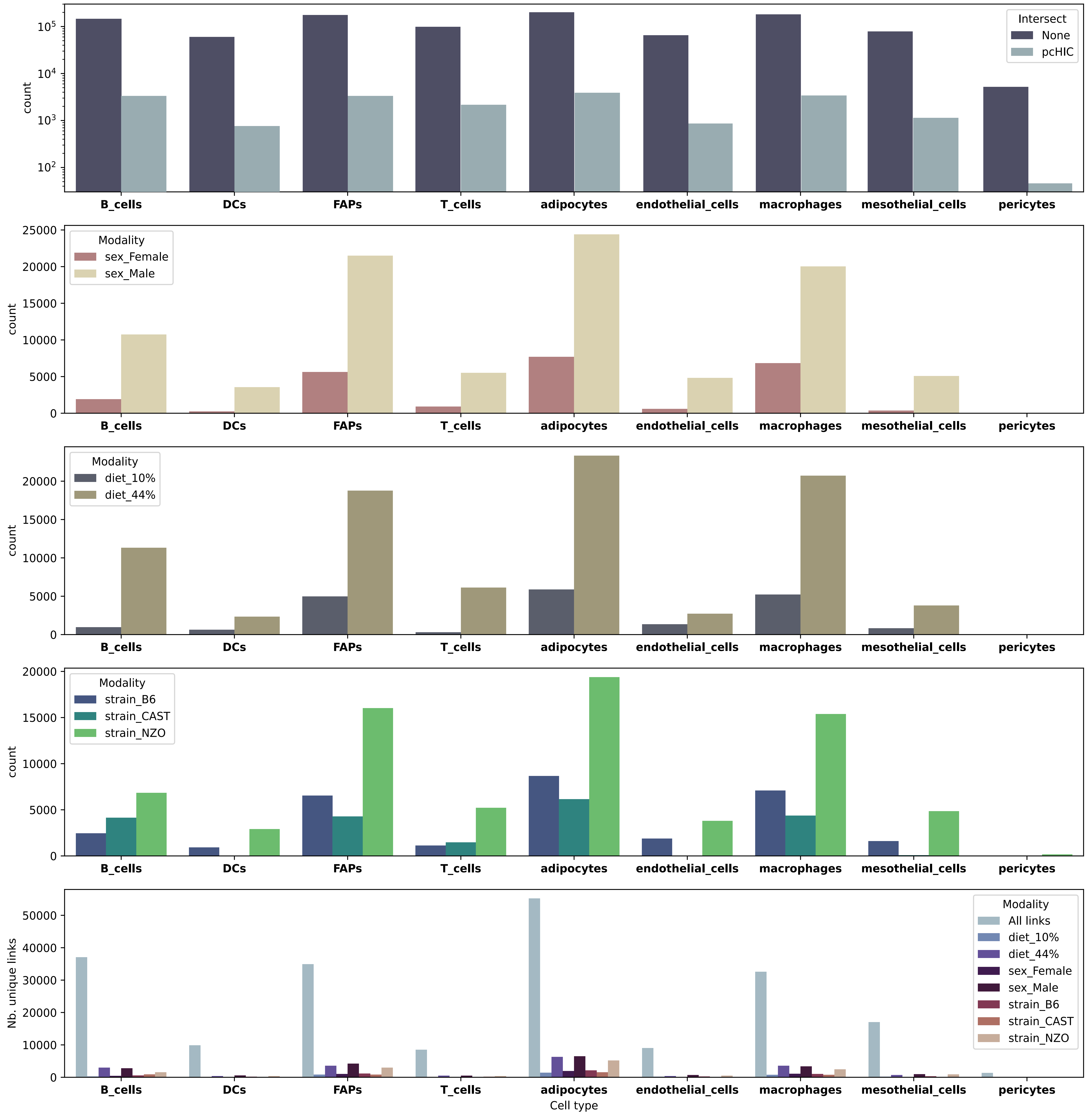

### Figure S7

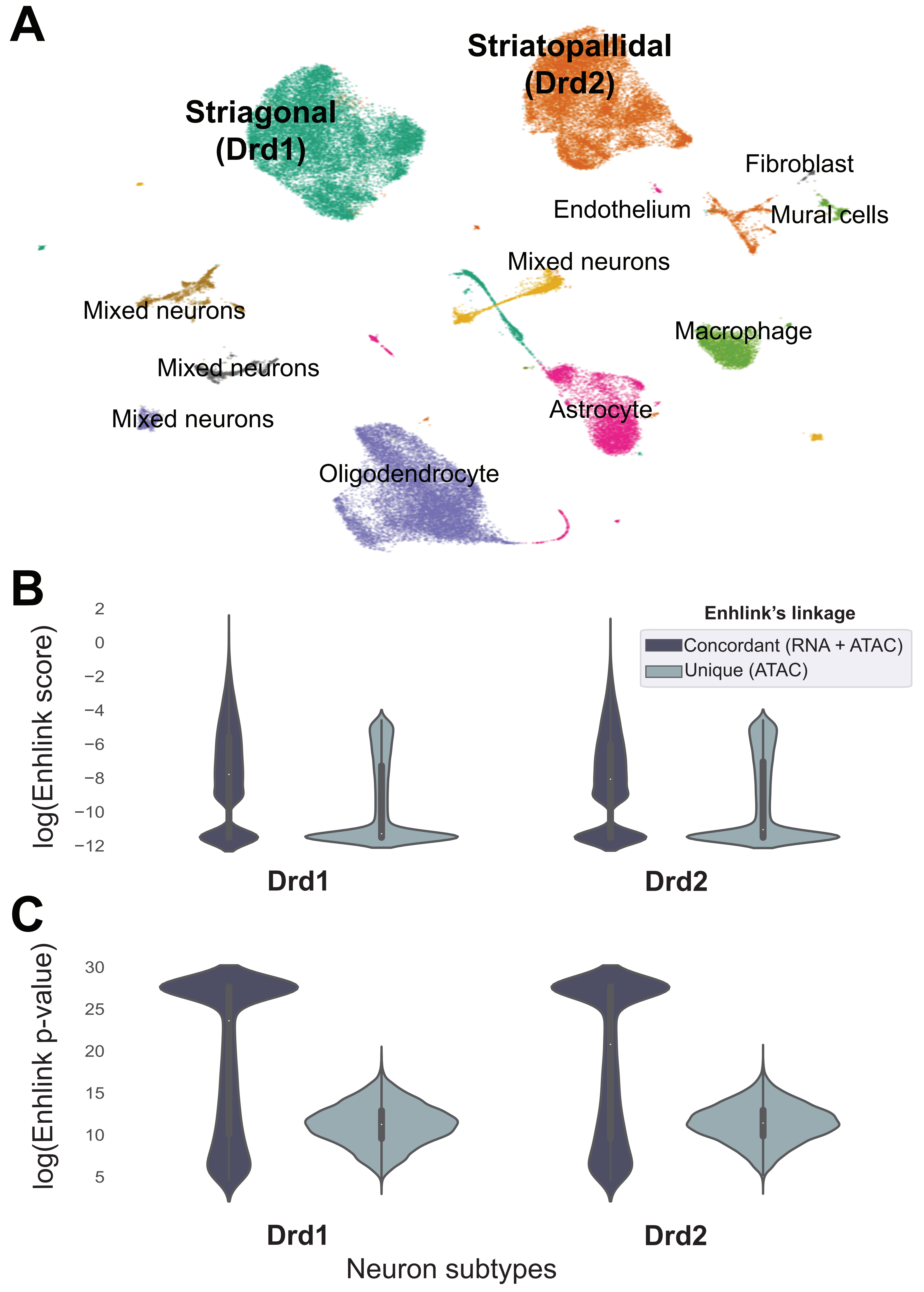

### Figure S8

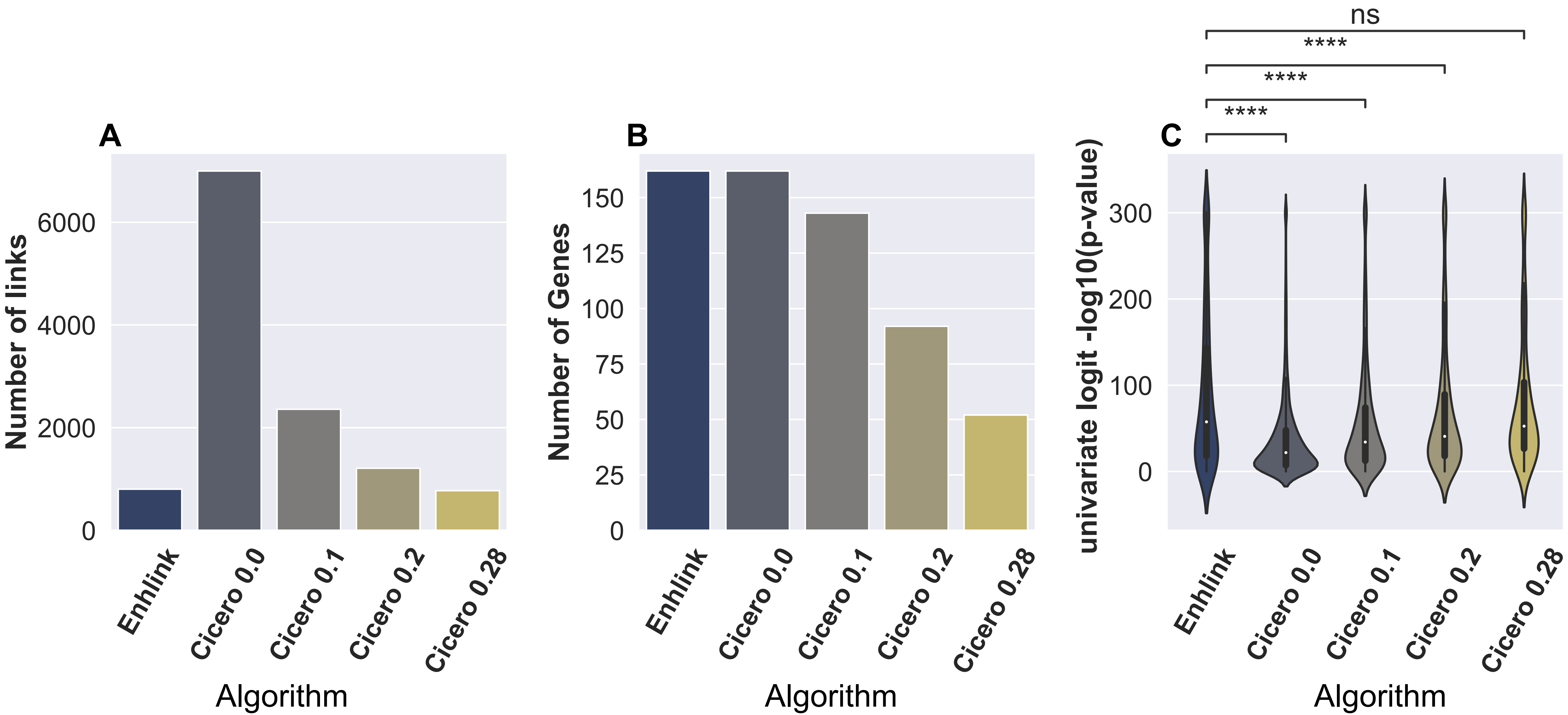

### Figure S9

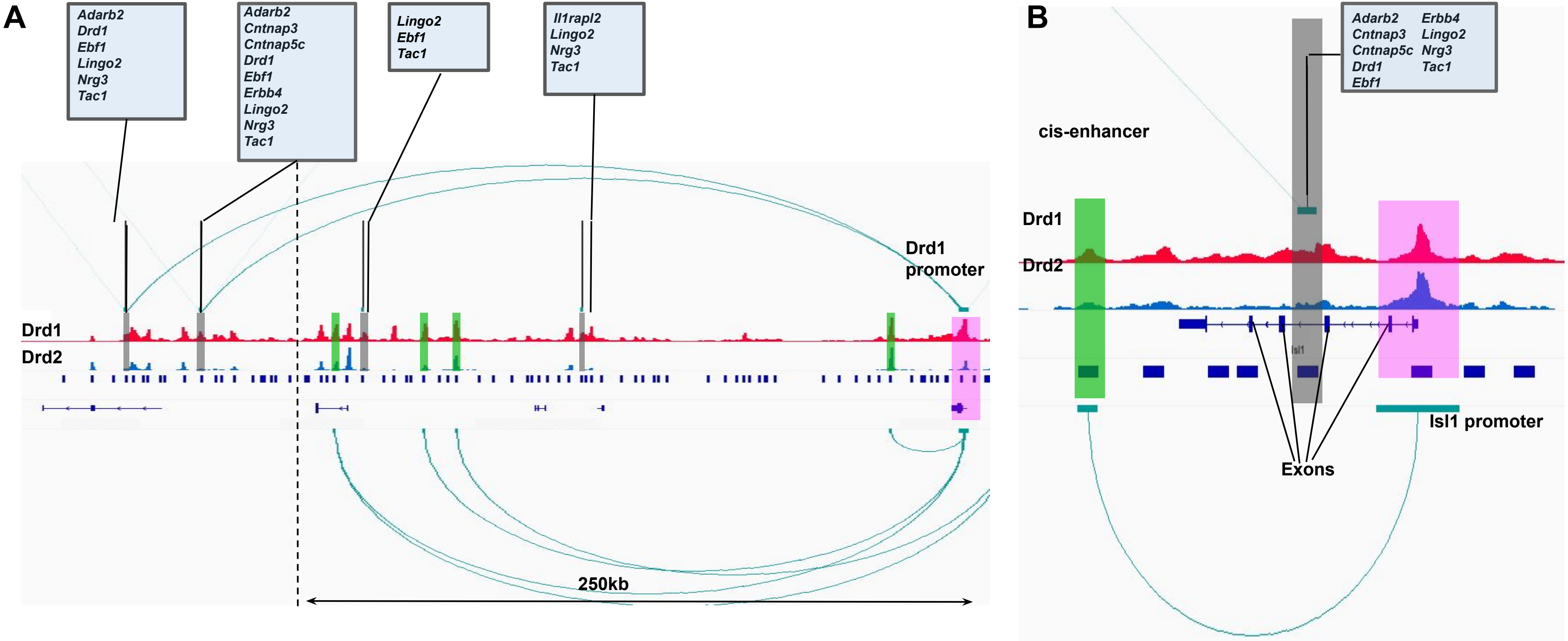
